# A Comprehensive Analysis of 3’UTRs in *Caenorhabditis elegans*

**DOI:** 10.1101/2024.02.15.580525

**Authors:** Emma Murari, Dalton Meadows, Nicholas Cuda, Marco Mangone

## Abstract

3’Untranslated Regions (3’UTRs) are essential portions of genes containing elements necessary for pre-mRNA 3’end processing and are involved in post-transcriptional gene regulation. Despite their importance, they remain poorly characterized in eukaryotes. Here, we have used a multi-pronged approach to extract and curate 3’UTR data from 11,533 publicly available datasets, corresponding to the entire collection of *C. elegans* transcriptomes stored in the NCBI repository from 2009 to 2023, and present its complete 3’UTRome dataset sequenced at single-base resolution.

This updated *C. elegans* 3’UTRome is the most comprehensive resource in any metazoan, covering 97.4% of the 20,362 experimentally validated protein-coding genes with refined and updated 3’UTR boundaries for 23,489 3’UTR isoforms.

We also used this novel dataset to identify and characterize sequence elements involved in pre-mRNA 3’end processing and update miRNA target predictions. This resource provides important insights into the 3’UTR formation, function, and regulation in eukaryotes.

## Introduction

The 3’ UnTranslated Regions (3’UTRs) are important segments within the gene structure located between the STOP codon and the poly(A) tail in eukaryotic mRNAs. They house sequences that serve as targets for various regulatory molecules, such as RNA binding proteins (RBPs) and small non-coding RNAs (ncRNAs), that exert control over gene expression by impacting mRNA biogenesis, translation rate, stability, and localization ^1,2^. During the mRNA transcription termination step, eukaryotic pre-mRNAs are cleaved by a large multi-subunit complex named the Cleavage and Polyadenylation Complex (CPC), which defines the length of 3’UTRs. The CPC identifies specific sequence elements within pre-mRNAs’ 3’UTRs and executes the cleavage reaction at the polyadenylation site (PS) element. The eukaryotic CPC contains at least 17 different subunits divided into four main subcomplexes; the Cleavage and Polyadenylation Specificity Factor (CPSF), the Cleavage stimulation Factor (CstF), and the Cleavage Factor 1 (CFIm) and 2 (CFIIm) (**Main Figure 1A**). The CPSF, CstF, and CFIIm subcomplexes, together with RRBP6 and Symplekin, are required for efficient 3’ end processing *in vitro* ^3,4^. In humans, the CPSF sub-complex is composed of the proteins CPSF1 (also known as CPSF160)^5,6^, CPSF2 (also known as CPSF100)^7^, CPSF3 (also known as CPSF73) ^7^, CPSF4 (also known as CPSF30)^5,6^, FIP1L1 ^8^ and WDR33 ^5,6^. CPSF4 and WDR33 recruit the CPSF subcomplex to the polyadenylation signal (PAS) element, a hexameric motif located ∼19 nucleotides upstream of the PS site ^5,9^, which in metazoans is canonically the sequence ‘AAUAAA’ ^10,11^, although several permutations of this element are sometimes used ^10,12,13^ (**Main Figure 1A**). Our group has recently identified a novel RRYRRR motif that can function in the absence of the canonical AAUAAA element in *C. elegans* ^14^. Following the docking of the CPSF subcomplex to the PAS element, the endonuclease CPSF3 then executes the cleavage reaction, and a string of adenosines is added at the 3’ end of the sequence by a specialized nuclear poly(A) polymerase, which in humans is named PAPOLA ^15^. FIP1L1 binds to U-rich sequences located between the PAS and the PS element ^8^ in an area named the buffer region ^14^ and stimulates the polyadenylation reaction ^8^ (**Main Figure 1A**).

**Main Figure 1:**
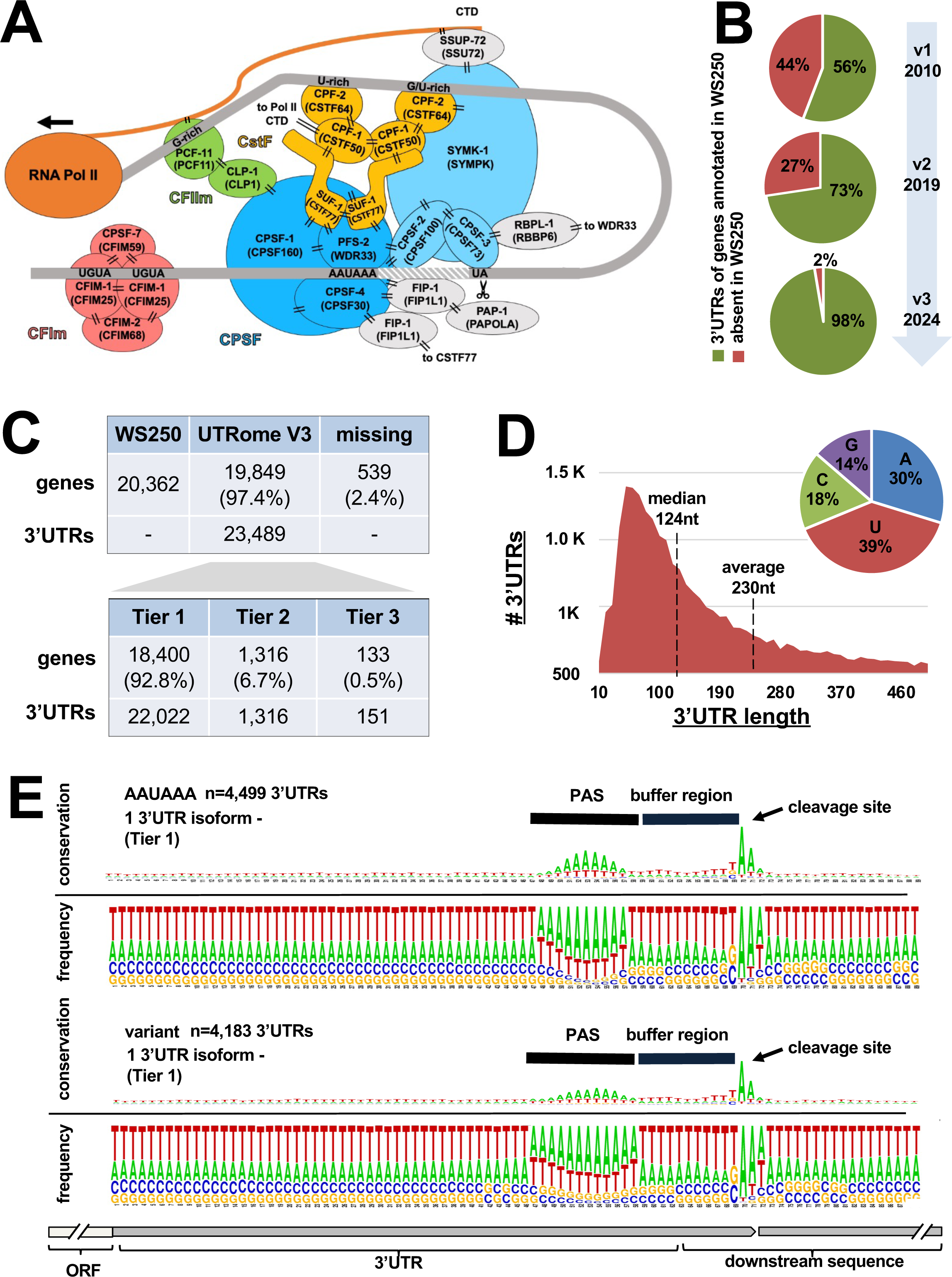
An overview of the *C. elegans* 3’UTRome. (A) The Cleavage and Polyadenylation Complex (CPC). Each member of this complex is labeled with the names of its *C. elegans* (top) and human (bottom) ortholog. Experimentally validated interactions between members are shown with two black dashes. A model 3’UTR is depicted in dark grey, the buffer region between the PAS and PS elements is highlighted with white stripes, and other elements involved in cleavage and polyadenylation are depicted in black text on the model 3’UTR. (B) Comparison between the updated 3’UTRome developed in this study (v3) and previously published versions ^10,14^. (C) (top) Overview of 3’UTR isoforms identified in this study. (bottom) Breakdown of 3’UTR isoform ranking. (D) Length distribution of the *C. elegans* 3’UTRome (v3). (inset) Nucleotide composition of the *C. elegans* 3’UTRome (v3). (E) Nucleotide conservation and frequency in genomic regions corresponding to 3’UTRs for genes with one 3’UTR isoform using a canonical (top) or a variant (bottom) PAS elements.

The CstF subcomplex is also needed for the mRNA cleavage reaction, but its role is less characterized. It is composed of a dimer of heterotrimers containing CSTF1 (also known as CSTF50), CSTF2 (also known as CSTF64), and CSTF3 (also known as CSTF77)^16^. In humans, the CSTF subcomplex binds to a pair of downstream sequence elements (DSEs), which are short motifs rich in guanine and uracil nucleotides (G/U-rich) located between 7-50 nucleotides downstream of the cleavage site ^16,17^ (**Main Figure 1A**). Lastly, the CFIIm subcomplex, which is the least characterized, consists of PCF11 and CLP1 ^18,19^. In humans, the CFIIm subcomplex binds to G-rich elements downstream of the CstF binding site ^19^ and is needed for the transcription termination reaction ^3^ (**Main Figure 1A**).

Studies in human cells also identified an additional upstream ‘UGUA’ sequence element recognized by the CFIm subcomplex that is not always present and is not required for the mRNA cleavage process *per se* but, when present, can act as a cleavage enhancer in the context of alternative polyadenylation ^20^ (**Main Figure 1A**). Importantly, the CPC may recognize different PAS elements within the same 3’UTR, producing transcripts with different 3’UTR lengths through a process known as alternative polyadenylation (APA) ^21^. Although the exact function of this process is unknown, it has been shown that in some cases, the shortening of 3’UTRs through APA allows genes to evade post-transcriptional gene regulation through the elimination of specific elements responsible for regulating mRNA processing and translation ^22–24^. How the CPC discriminates between PAS elements within the same 3’UTR is also unclear. Still, disturbances in the 3’ end processing of pre-mRNAs have far-reaching effects on diverse developmental and metabolic processes, and they are linked to the emergence and advancement of many diseases, including neurodegenerative disorders, diabetes, and cancer ^21,23^.

Despite their importance, 3’UTRs are poorly annotated in all metazoans. While a human comprehensive 3’UTR dataset is still not available, several efforts have been made in the past years to complete 3’UTR datasets in model organisms, including *C. elegans* ^10,12,14^ and *D. melanogaster* ^25^. Although not saturated, these datasets provided an overview of 3’UTR formation and their regulation. Additionally, since the functional domains and key amino acids of the members of the CPC are highly conserved across metazoans ^14^, investigating 3’UTR formation and how it influences 3’UTR function in model systems provided valuable insights into how these processes function in other metazoans, including humans.

Throughout the years, our lab has refined the 3’UTR collection in *C. elegans* in the whole animal and in several somatic tissues in various stages of development^10,13,14,24,26,27^ (**Main Figure 1B**).

Our previous *C. elegans* 3’UTRome (v2), which is currently available in WormBase, contains 3’UTR boundaries for 14,788 genes corresponding to 23,084 3′UTR isoforms (73% of all protein-coding genes included in the WS250 release) (**Main Figure 1B**).This dataset was not yet saturated since we could not assign 3’UTR data to the remaining 5,554 protein-coding genes in WS250 (**Main Figure 1B**).

Here, we have used a multi-pronged approach consisting of 1) remapping 3’UTR data from the entire collection of *C. elegans* transcriptomes stored in the NCBI Sequence Read Archive (SRA) repository from 2009 and 2023 (11,533 transcriptome datasets; 240TB in size), 2) incorporating data from past 3’UTR datasets ^10,12,14^, and 3) adding a high-throughput cloning pipeline to detect the presence of rare 3’UTRs. We have also used this resource to study the 3’UTR elements involved in their biogenesis and function, including the PAS element, PS element, the buffer region, and updated obsolete miRNA target predictions. Our approach produced the most comprehensive 3’UTR resource in any metazoan to date, covering ∼98% of the *C. elegans* transcriptome with an ultra-deep coverage.

## Results

### A complete *C. elegans* 3’UTRome dataset

Our updated *C. elegans* 3’UTRome (v3) was prepared using three major pipelines (Tiers) (**Main Figure 1C** and **Supplemental Figure S1)**. The first two Tiers (1 and 2) were prepared by extracting 3’UTR boundary information from 11,533 *C. elegans* transcriptome datasets from the SRA repository (**Supplemental Table S1**). We reasoned that this large amount of data contains 3’UTRs from all possible cleavage sites recognized by the CPC in this model system. The first one (Tier 1) is the strictest and contains 3’UTR data for 93% of the protein-coding genes present in this dataset (18k genes, 22k 3’UTR isoforms) (**Main Figure 1C**). Tier 2 was prepared by lowering our filter threshold to identify rare 3’UTRs not detected in our stringent Tier 1 dataset. This pool is small and contains ∼ 7% of the mapped 3’UTRs. Lastly, Tier 3 includes 3’UTRs from past 3’UTRomes ^10,12,14^. Additionally, we developed a medium-throughput cloning pipeline targeting rare 3’UTR isoforms not present in our Tiers. (**Supplemental Figure S2**). Taken together, this new resource contains 3’UTR data for 98% of all *C. elegans* protein-coding genes (**Main Figure 1C and Supplemental Table S2**).

The 3’UTRs in the *C. elegans* have similar properties to those characterized in earlier studies ^10,12,14^. They have a median length of 124 nucleotides and are rich in adenosine and uracil nucleotides (**Main Figure 1D**). Notably, aligning these 3’UTRs at the cleavage site reveals a strong enrichment of adenosines, with some enrichment of uracil nucleotides at the second position from the cleavage site and a buffer region length of ∼14 nucleotides (**Main Figure 1E**)

### The PAS element in *C. elegans* uses an RRYRRR motif

Next, we used the Tier 1 3’UTRs from our 3’UTRome to further analyze PAS element usage and the prevalence of APA in *C. elegans.* As predicted, most 3’UTRs use the canonical PAS element AAUAAA (47%) (**Main Figure 2A**). Notably, we observed increased occurrences of variant PAS usage in comparison to earlier 3’UTRome datasets (53% of the PAS elements detected in v3 vs 42% detected in v2) ^14^ (**Main Figure 2A**). 68% of these variant PAS elements maintain the chemical nature of the canonical PAS site (RRYRRR). We also frequently observe the occurrence of this RRYRRR motif where the PAS element is usually located (**Main Figure 2B**). Strikingly, ∼99% of all PAS identified in this study contain a Y3 and R6 nucleotide within the PAS element, remarking the importance of the relative location and the chemical nature of these two nucleotides in the PAS sequence, as previously shown using CryoEM data^5,6^. We have listed all the variant PAS elements permutations and the gene data in **Supplemental Table S3**.

**Main Figure 2:**
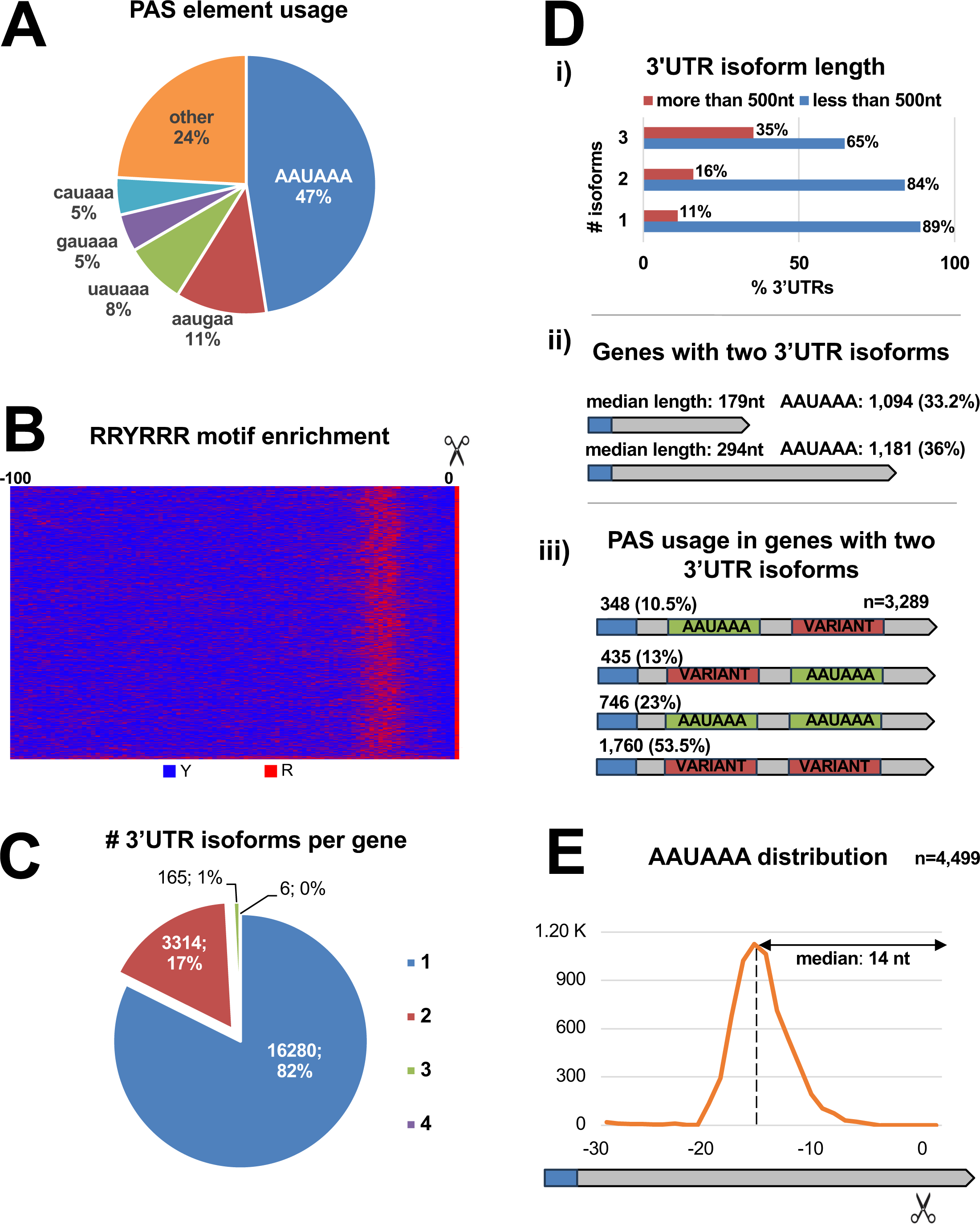
PAS element usage in the *C. elegans* 3’UTRome v3. (A) PAS element usage. The top 5 PAS elements are shown. (B) Pyrimidine (blue) and purine (red) distribution within the terminal 100 nt of 3’UTRs from protein-coding genes with one 3’UTR isoform. (C) Number of genes with multiple 3’UTR isoforms identified in this study. (D) PAS usage and 3’UTR length in genes with multiple 3’UTR isoforms. i) Proportions of genes with 3’UTR isoforms greater or less than 500 nucleotides in length separated by the number of 3’UTR isoforms. ii) Median length and canonical PAS element usage of proximal and distal 3’UTR isoforms in genes with two 3’UTR isoforms. iii) Occurrence of canonical (green) and variant (red) PAS usage in genes with two 3’UTR isoforms. (E) Distribution of canonical PAS element location in 3’UTRs from genes with one 3’UTR isoform.

### Alternative polyadenylation occurs but is uncommon in *C. elegans*

82% of *C. elegans* protein-coding genes use only one 3’UTR isoform. 17% of genes use two 3’UTR isoforms, and only 1% of genes have three or more 3’UTR isoforms, indicating that APA in this model system is less extensive than what was previously found ^14^ (**Main Figure 2C**). Genes with increased 3’UTR isoforms are more likely to have longer 3’UTRs (**Main Figure 2D top panel**). The median length difference between 3’UTRs with two isoforms is 115 nucleotides (**Main Figure 2D middle panel**). Interestingly, the median length of the short 3’UTR isoform in genes with two 3’UTR isoforms is 179 nucleotides, longer than the median 3’UTRs identified in this study (**Main Figure 1C**). The increased length in 3’UTRs that use APA suggests that there may be more elements not yet characterized involved in this process in *C. elegans*. Notably, many of the distal 3’UTR isoforms of these alternatively polyadenylated genes are targeted by miRNAs that do not target the proximal 3’UTR isoform, suggesting each 3’UTR isoform is subject to different degrees of regulation (**Supplemental Figure S3)**.

Variant PAS are commonly present in genes with multiple 3’UTR isoforms (**Main Figure 2D bottom panel**). Unexpectedly, the canonical AAUAAA element is the second most abundant PAS element in genes with multiple 3’UTR isoforms (23%) (**Main Figure 2D bottom panel**), suggesting a potential novel regulatory mechanism in place used by the CPC to discriminate between similar PAS elements within the same 3’UTR.

### The nucleotide composition of the buffer region does not impact 3’UTR processing

The median length of the *C. elegans* buffer region is 14 nucleotides (**Main Figure 2E**), and is the same for canonical and variant PAS-containing 3’UTRs (*data not shown*). Nucleotide conservation analysis of buffer regions of various lengths from Tier 1 3’UTRs using canonical PAS elements showed a slight enrichment of uracil nucleotides at the fourth and fifth nucleotides downstream from the PAS element (**Main Figure 3A, yellow box**). This enrichment is independent of the length of the 3’UTR. We speculate that this region might be recognized by FIPP-1 and used by this factor to stabilize the CPC, though further investigation is needed to confirm this hypothesis.

**Main Figure 3:**
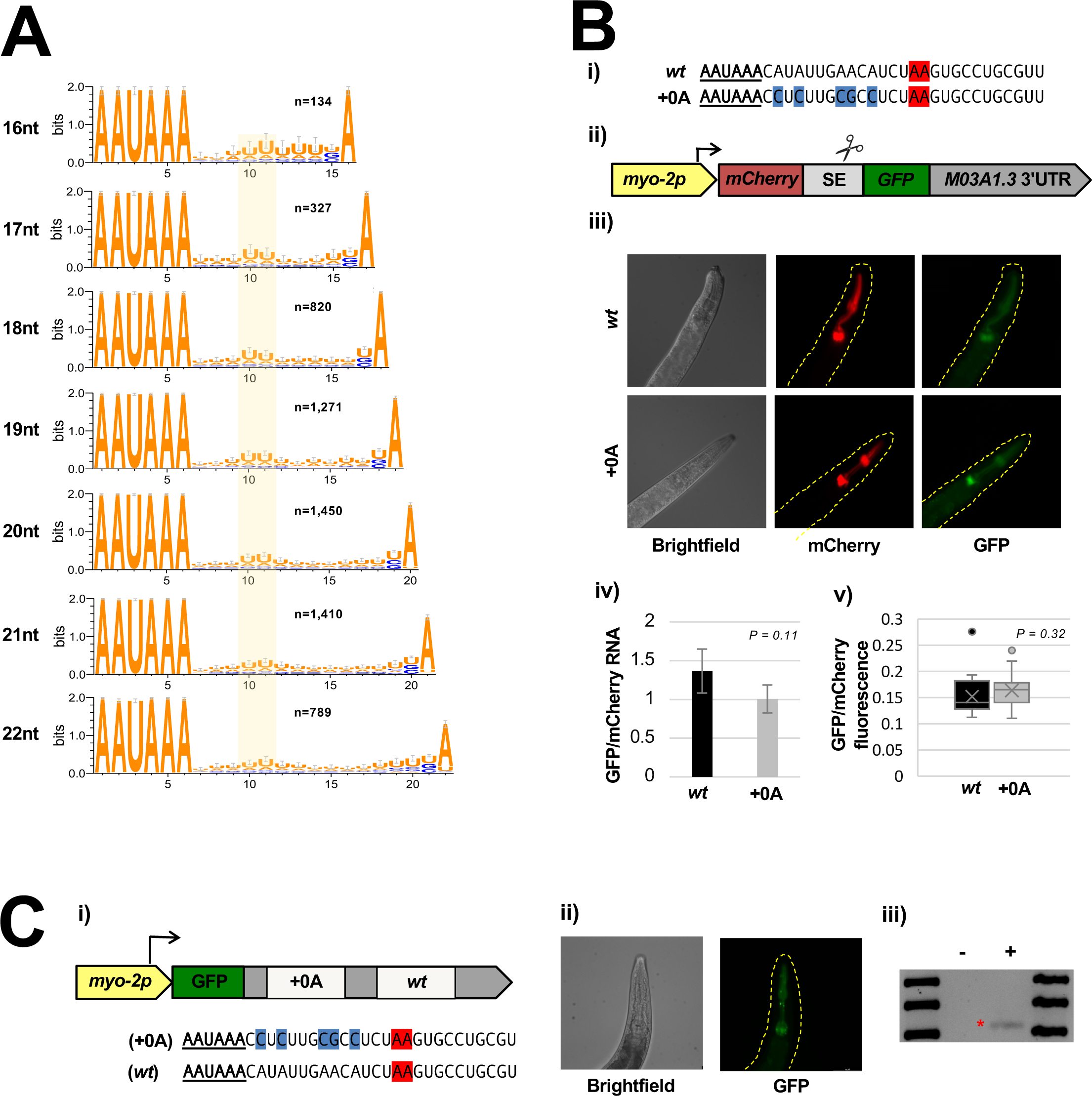
Effect of nucleotide composition of the buffer region on the cleavage and polyadenylation reaction. (A). Logo plot showing nucleotide conservation in buffer regions of various lengths from 3’UTRs from protein-coding genes with one 3’UTR isoform and a canonical PAS element. An enrichment of uracil nucleotides at +4 nucleotides from the PAS element is highlighted in yellow. (B) (i). The *wt M03A1.3* 3’UTR (top) and the +0A mutant *M03A1.3* 3’UTR, which lacks adenosines in the buffer region (blue) (bottom). The predicted cleavage site is highlighted in red. The canonical PAS element is underlined. (ii) A model of the dual fluorochrome constructs used to express the test 3’UTR *M03A1.3* (grey). mCherry (red) and GFP (green) are connected by a spliceable element (SE). The construct is expressed in the pharynx muscle using the *myo-2* promoter (yellow). (iii). Brightfield (left), mCherry (center), and GFP (right) images of representative animals expressing the *wt* and +0A dual fluorochrome transgenes. (iv). Ratio of mCherry and GFP mRNA levels, as determined by qPCR, between *wt* and +0A strains. (v). Ratio of mCherry and GFP fluorescence levels between the *wt* and +0A strains. (C). (i). A model of the construct used to test the buffer region’s influence on the cleavage site decision. Two sites (+0A and *wt*) were cloned into the test 3’UTR fused to a GFP reporter (green) and expressed in the pharynx muscle using the *myo-2* promoter (yellow). The sequences of the two polyadenylation sites are shown below the model. (ii). Representative brightfield (left) and GFP (right) images of the transgenic line of *C. elegans* expressing the construct. (iii). RT-PCR analysis of PAS usage with (+) or without (-) reverse transcriptase enzyme. The red asterisk highlights the presence of the +0A isoform.

In previous experiments, we observed a potential structural role for the buffer region in the cleavage and polyadenylation reaction ^14^. In those experiments, changing the length of the buffer region in several test 3’UTRs *in vivo* altered the location of their respective PS sites, suggesting that perhaps the buffer region may fold or assume secondary structures incompatible with the cleavage reaction ^14^. To test this hypothesis, we removed all five adenosine nucleotides in the buffer region in one of these test 3’UTRs (*M03A1.3)* to disrupt its nucleotide composition and potentially its secondary structure.

We used an *in vivo* dual-color reporter system that we developed in the past ^13,24^. This system drives the expression in the pharyngeal tissue of two fluorochromes (GFP and mCherry) separated by a spliceable element. Upon transcription, these two fluorochromes are translated independently; the mCherry expression reports transcription levels and the GFP expression reports translation levels.

We generated two *C. elegans* strains by cloning the *wt* M03A1.3 3’UTR and a mutant 3’UTR lacking five adenosine nucleotides (+0A) (**Main Figure 3B**). Both strains express similar ratios of mCherry and GFP (*p=0.32*). This result is also echoed by qPCR analysis, showing no significant difference between these two strains (p=0.11) (**Main Figure 3B**). We also cloned both +0A and the *wt* buffer regions in tandem within a test 3’UTR fused to GFP fluorochrome and used RT-PCR experiments to test if the +0A buffer region can dominate over the wt buffer region in executing the cleavage and polyadenylation reaction. These results showed that the +0A buffer region is sufficient to direct cleavage at the correct site (**Main Figure 3C**). Taken together, these experiments suggest that altering the nucleotide composition of the buffer region does not impact gene expression or polyadenylation site usage.

### The adenosine nucleotide in the PS site defines the end of the 3’UTR

In previous experiments, we identified an enrichment of adenosine nucleotides at the cleavage site ^14^. This enrichment was functional since its removal altered, in some cases, the location of the PS site ^14^. However, the exact function of this terminal adenosine remained poorly understood.

In our new dataset, 89% of *C. elegans* 3’UTRs terminate with an adenosine nucleotide, suggesting a role for this specific nucleotide in determining PS site location (**Main Figure 4A**). To further study its role *in vivo*, we generated five scanning insertion mutants in the 3’UTR of the gene *M03A1.3*, extending the buffer region up to an additional eight nucleotides (+0A, +2A, +4A, +6A, or +8A). We then prepared transgenic *C. elegans* strains and used them to test if each of these added adenosine nucleotides could relocate the position of the PS site independently of the PAS element (**Main Figure 4B**). As expected, in the +0A mutant, the cleavage always occurs at the canonical site (n=30). Strikingly, in the +2A mutants, the cleavage reaction shifted downstream at the newly inserted adenosines even if the PAS location was the same (62%), suggesting that terminal adenosine can indeed dominate over the PAS site in determining the PS site location (n=13). In the +4A, +6A, and +8A mutants, we observed little to no cleavage at the newly added terminal adenosine (**Main Figure 4C**), perhaps because the PAS element and the PS element in these mutants are too far apart for the CPC to process them correctly.

**Main Figure 4:**
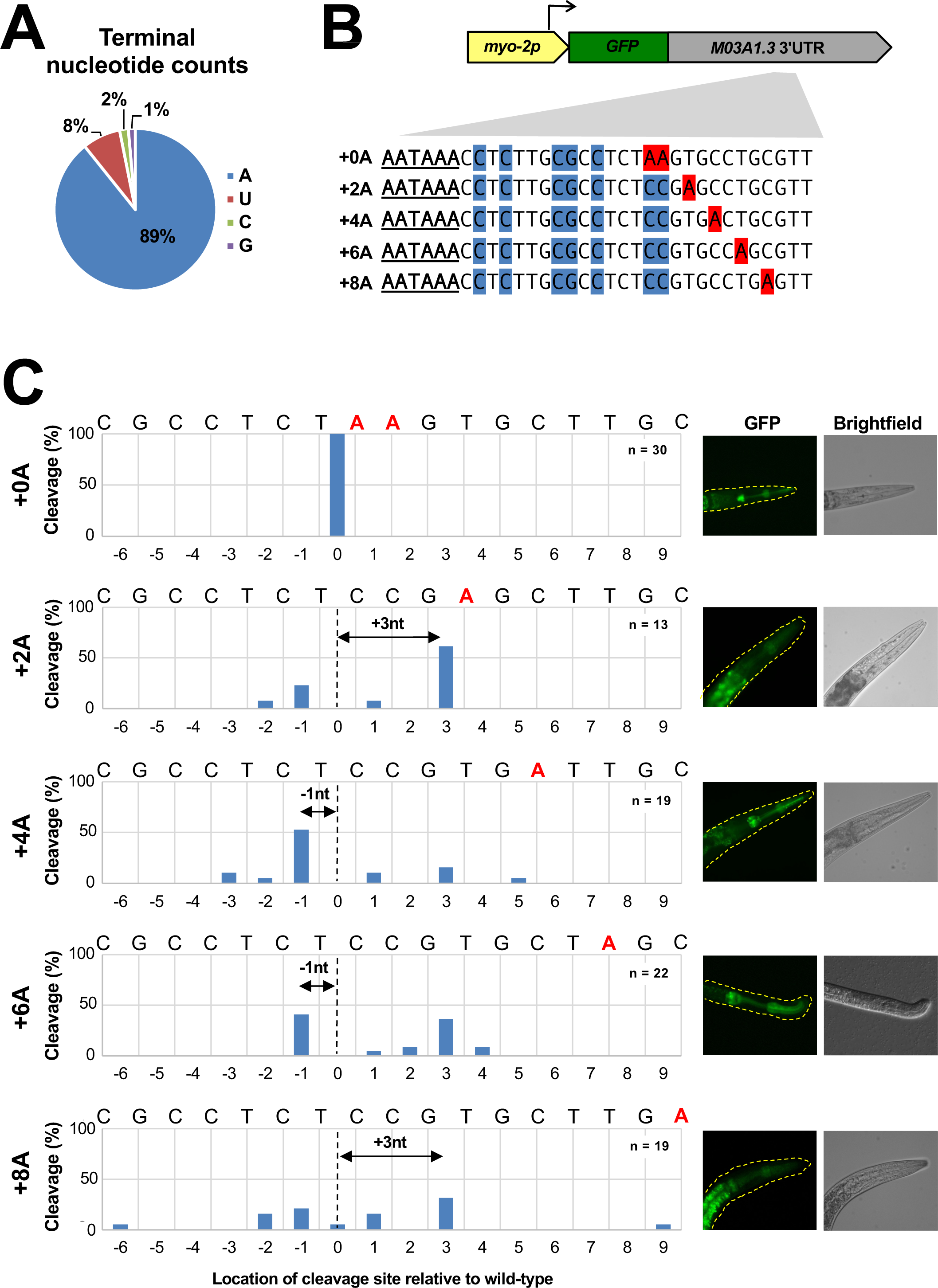
The terminal adenosine influences the location of the cleavage site. (A). Terminal nucleotide counts of the 3’UTRs in the *C. elegans* 3’UTRome (v3). (B). Scanning insertion mutant analysis. Adenosine nucleotides in the buffer region and at the cleavage site in the *M03A1.3* test 3’UTR were replaced with cytosines and guanines (blue). Adenosine nucleotides were inserted +2, +4, +6, and +8 nucleotides downstream of the canonical cleavage site (+0A) (red). The canonical PAS element is underlined. Each *M03A1.3* 3’UTR mutant was cloned into a construct containing GFP (green) and the *myo-2* promoter (yellow). (C). Distribution of the location of the cleavage reaction in transgenic animals expressing the scanning insertion constructs. The adenosines at the predicted cleavage sites are highlighted in red. The dotted line marks the *wt* cleavage site. The arrows mark the distance between the most used cleavage site in each mutant and the *wt* cleavage site. Representative GFP (left) and Brightfield (right) images of each mutant are shown to the right of each bar chart.

In addition to their inability to define new PS sites, most of the +4A and +6A mutants were cleaved at a new cryptic site between a cytosine and an uracil located one nucleotide upstream of the canonical cleavage site. Additionally, most of the +8A mutants and many of the +6A mutants were cleaved at another cryptic cleavage site located between a guanine and an uracil nucleotide three nucleotides downstream of the canonical cleavage site (**Main Figure 4C**).

Taken together, our results suggest a direct role for terminal adenosines in determining the PS site, which is lost if the buffer region becomes too long. In this case, new cryptic sites are used instead.

### The uracil nucleotide preceding the terminal adenosine does not influence the location of the cleavage reaction

Most of the novel cryptic sites in our scanning deletion analysis occur at an uracil nucleotide lacking a terminal adenosine nucleotide, suggesting that this uracil can substitute it at the PS site. (**Main Figure 4C**). Strikingly, 49% of 3’UTRs in this new *C. elegans* 3’UTRome (v3) possess an uracil nucleotide at −1 from the PS site, next to a terminal adenosine nucleotide **(Main Figure 5A**), suggesting a possible role for this nucleotide in PS recognition. To test this hypothesis, we repeated our scanning insertion assay by preparing a new mutant, replacing the uracil preceding the terminal adenosines with a guanine nucleotide (+0GA). We hypothesized that if an uracil nucleotide was needed at this position, this new mutant would disrupt the PS site decision or at least make it more variable. Instead, we notice that in the absence of this uracil nucleotide, the cleavage still occurs precisely at the original PS site (**Main Figure 5C, Top panel**). To validate this result, we replaced a guanine preceding the terminal adenosine in our +2A mutant background with an uracil nucleotide (+2TA). Also, with this mutant, the cleavage location was not disrupted (**Main Figure 5C, Bottom panel**). Taken together, these results suggest that this uracil nucleotide does not influence the exact location of the cleavage site per se. Still, if present, it may play a different role in the cleavage reaction, such as increasing the likelihood of successful cleavage (**Main Figure 5C**). However, more investigation is needed to confirm this.

**Main Figure 5:**
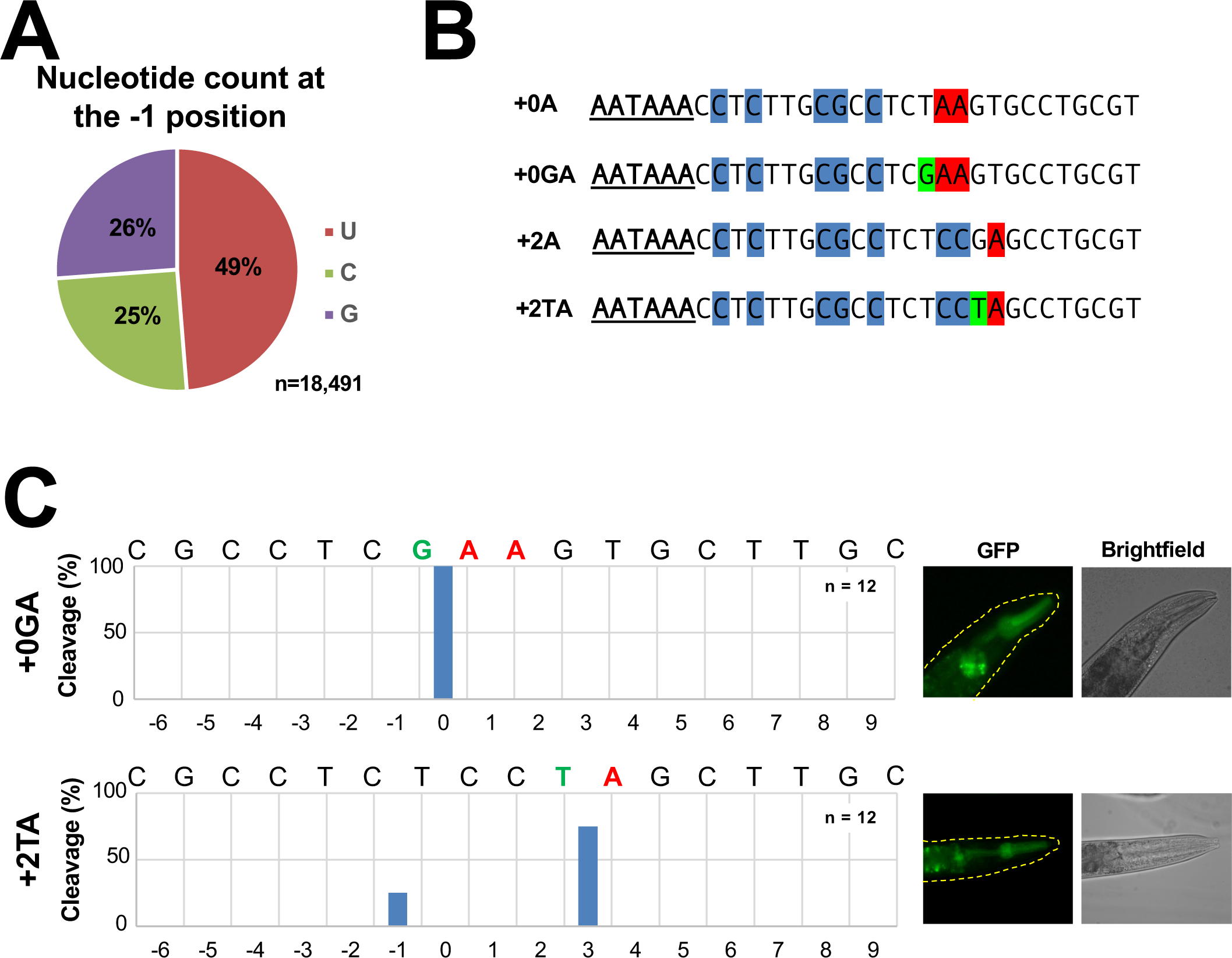
The nucleotide preceding the terminal adenosine does not influence the location of the cleavage site. (A) Nucleotide usage at −1 from the cleavage site in the *C. elegans* 3’UTRome (v3). (B) Sequences of the +0GA and +2TA mutants. These mutants resemble the +0A and +2A mutants, but the nucleotides in the −1 position have been replaced (green). (C) Distribution of the location of the cleavage reaction in transgenic animals expressing +0GA (top) and +2TA (bottom) constructs. The adenosines at the predicted cleavage sites are highlighted in red, and the −1 position nucleotides are highlighted in green. Representative GFP (left) and Brightfield (right) images of each mutant are shown to the right of each bar chart.

### *C. elegans* PAS elements are not always located in 3’UTRs

Notably, in the +8A mutant tested in **Main Figure 4C**, we identified one novel cryptic cleavage site that uses a curious variant PAS site (UAUAAA) located 12 nt upstream of the STOP codon (**Supplemental Figure S4**). Only 2 of the 21 sequenced +8A mutant transcripts used this cryptic PAS site, indicating that its usage is rare.

In our 3’UTRome (v3), 107 protein-coding genes naturally possess a PAS site upstream of a STOP codon (**Supplemental Figure S5A**). These ORF-PAS 3’UTRs are relatively short, with most containing only ∼11 nucleotides (**Supplemental Figure S5B**). Half use a canonical AAUAAA PAS element (**Supplemental Figure S5C**). The complete list is shown in **Supplemental Table S4**.

### *C. elegans* possesses U-rich DSEs

The *C. elegans* downstream sequence elements (DSEs) contain binding sites for CstF and are poorly characterized. To identify these elements in our 3’UTRome v3, we extracted and analyzed genomic regions from the cleavage site to 100 nt downstream of it. These regions contain an enrichment of uracil nucleotides located between +1 and +20 nucleotides downstream of the cleavage site (**Supplemental Figure S6, Top panel**). MEME analysis of the 100 nucleotide genomic regions revealed a symmetrical bipartite U-rich element that could represent a potential binding site for the CstF subcomplex in *C. elegans* (**Supplemental Figure S5, bottom panel**). This analysis did not reveal any G/U-rich DSEs, suggesting the *C. elegans* CstF subcomplex may only bind to U-rich DSEs.

### The upstream UGUA element is not present in *C. elegans* 3’UTRs

Some human 3’UTRs contain upstream elements with the sequence ‘UGUA’ that act as cleavage enhancers ^20^. We attempted to detect these elements by extracting the last 100 nucleotides of distal and proximal 3’UTR isoforms of several *C. elegans* protein genes that use APA and are transcribed with two 3’UTR isoforms. There is no enrichment of the UGUA element in either the proximal or distal isoforms (**Supplemental Figure S7A**). Additionally, when present, this ‘UGUA’ motif is evenly distributed throughout *C. elegans* 3’UTRs (**Supplemental Figure S7B)**, indicating that *C. elegans* 3’UTRs may not use strict upstream sequence elements containing the ‘UGUA’ motif.

### The *C. elegans* 3’UTRome v3 allows for making more accurate miRNA targeting predictions

Since we now have 3’UTR coordinates for ∼98% of all *C. elegans* protein-coding genes, we use this information to update miRNA targeting predictions using the miRanda algorithm ^28^. We prepared three datasets; the first one (strict+) is very stringent, requires perfect miRNA seed complementarity to its target, and requires a binding energy score > 300. It contains 4,448 predictions for 2,469 protein-coding genes. The second dataset (strict) is also stringent but does not account for the binding energy score. It contains 140,640 predictions for 17,207 unique protein-coding genes. The third dataset (all) does not include seed complementarity or binding energy score requirement and contains 322,959 predictions for 18,701 protein-coding genes. We used the strict+ dataset to generate a network of all the miRNA-mRNA interactions in the *C. elegans* transcriptome (**Main Figure 6A**). The network is very dense, with 1,291 nodes and 1,666 edges. Known interactions, such as those between *let-7* and *hbl-1* ^29^, *let-7* and *daf-12* ^30^ (**Main Figure 6B**), and *lsy-6* and *cog-1* ^31^ are shown (**Main Figure 6C**). The most connected miRNA is *miR-247-5p*, which has predicted binding sites for 202 unique protein-coding genes, followed by *miR-85-5p*, with predicted binding sites for 184 unique protein-coding genes. The heterochronic regulator gene *lin-14* is instead the most connected gene in this network, having predicted binding sites in its 3’UTR for nine different miRNAs (*let-7-5p, lin-4-5p, miR-40-5p, miR-48-5p, miR-84-5p, miR-237-5p, miR-241-5p, miR-795-5p, and miR-794-5p*). *Lin-14* is followed by the egg-laying abnormal gene *egg-46*, which possesses seven predicted miRNA binding sites, and the hunchback-like gene *hbl-1*, with six. Unfortunately, using the strict+ dataset does not contain some known interactions, such as the interactions between *let-7* and *lin-41* ^32,33^, *let-7* and *lin-60* ^32,34^ and *lin-4* and *lin-28* ^35^ due to their low binding energy values. Because of this, we have included all three datasets in **Supplemental Table S5**.

**Main Figure 6:**
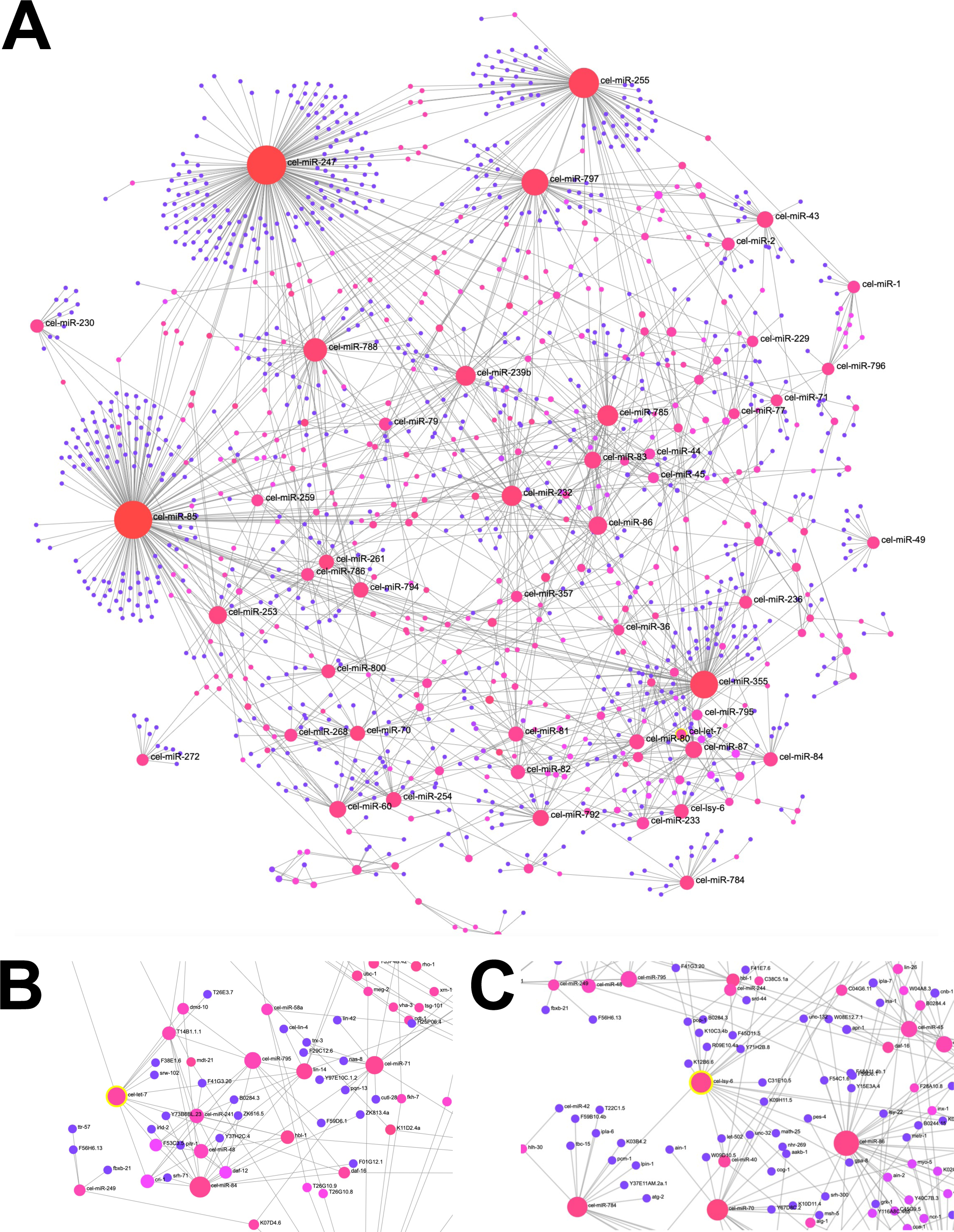
The *C. elegans* miRNA/3’UTR interactome. (A). Network showing miRNA-3’UTR interactions identified in this study. The miRNA interactions shown are included in the strict dataset (see **Supplemental Table S5**) miRNAs are depicted by dark pink dots. The size of each of these dots corresponds to the number of predicted interactions. Genes with one interaction are depicted by purple dots, and genes with multiple interactions are depicted by pink dots. (B). Network inset showing *let-7* and its predicted targets. (C). Network inset showing *lsy-6* and its predicted targets.

## Discussion

The mapping of all 3’UTRs in *C. elegans* protein-coding genes was initiated in 2010 (v1) under the modENCODE project, which provided 3’UTR data for 11,369 *C. elegans* protein-coding genes ^26^ (**Main Figure 1B**). This collaborative effort generated the first 3’UTRome for any metazoan organism. Although it only mapped 3’UTR isoforms for ∼56% of all protein-coding genes in WS250, it allowed these portions of genes to be studied at an unprecedented resolution. Another dataset produced later validated and improved this initial study ^12^. Ten years after the release of the v1 dataset, a second, more comprehensive dataset was released (v2) ^14^, which included 3’UTR data from both v1 and Jan *et al*. The v2 expanded both previous studies, mapping 3’UTR data for an additional ∼3,500 protein-coding genes and covering ∼73% of all protein-coding genes (**Main Figure 1B**).

In this study, our latest version of the *C. elegans* 3’UTRome (v3) contains 3’UTR isoform data for ∼98% of the genes in the WS250 release, making it the first complete 3’UTRome for any organism (**Main Figure 1B-C**). We obtained this information by extracting 3’UTR data from all the *C. elegans* transcriptome datasets stored in the SRA database from 2009 to 2023 and cloning and sequencing rare 3’UTRs that this analysis might have missed (**Supplemental Figure S2**). With this complete dataset, we can perform more accurate analyses to identify and characterize various regulatory and processing elements in 3’UTRs, including miRNA targets and those involved in pre-mRNA 3’end processing.

Importantly, we could not detect 3’UTR data for 513 protein-coding genes in WS250. Our targeted cloning approach described in **Supplemental Figure S2** failed to identify ∼70% of a test pool of 83 test genes for which we could design primers (**Supplemental Table S7**). This list is included in **Supplemental Table S8**. Some of these genes, such as histone genes, are known to lack a polyA tail, and may have escaped our cloning approach. Since we re-analyzed 11,533 datasets and did not map any 3’UTRs for those genes, we doubt that they exist, and they may be misannotated in WS250.

The PAS element is one of the processing elements located within eukaryotic 3’UTRs. It was discovered ∼50 years ago as the first element involved in mRNA cleavage and polyadenylation ^11,36,37^. In metazoans, it has the canonical sequence AAUAAA, though many other variant sequences are often used ^11^ **(Main Figure 2A).** Since many permutations of this element have been identified in the past, it is not clear how the CPC can identify, discriminate, and use these sometimes very different PAS sequences. Our lab recently showed that in the absence of the canonical “AAUAAA” element, the variant PAS sequence frequently contains an RRYRRR motif ^14^. This remains true in this updated worm 3’UTRome (v3), as ∼68% of PAS elements possess this RRYRRR motif **(Main Figure 2C).** Notably, most of these variant PAS elements contain a pyrimidine at the third position and a purine at the sixth position, indicating that the chemical nature of these nucleotides plays a particularly important role in PAS recognition. Previous studies of the human CPC provided mechanistic insights, showing that the uracil nucleotide at the third position and the adenosine nucleotide at the sixth position in the canonical “AAUAAA” PAS element base pair with one another to form a structure that fits within part of the CPSF subcomplex during the cleavage and polyadenylation reaction ^5,6^. These nucleotides likely play a similar role during cleavage and polyadenylation in *C. elegans.* Surprisingly, 99% of all PAS elements identified in this study possess a pyrimidine in 3^rd^ position and a purine in 6^th^ position. Also, the remaining 1% still possess these nucleotides but in reverse order, indicating that these nucleotides’ ability to base pair with one another is the chemical property most essential for PAS element function. The location of the PAS element is also important. In *C. elegans* all protein-coding genes possess this element between −20nt and −9nt from the PS site, with a median distance of –16nt **(Main Figure 2E**).

18% of *C. elegans* protein-coding genes use alternative polyadenylation (**Main Figure 2C and Supplemental Figure S3**). This apparent difference from our earlier report (42%) ^14^ is mostly due to our stringent algorithm, which assigns 3’UTR clusters to protein-coding genes. 3’UTRs in protein-coding genes utilizing APA tend to have a longer average length **(Main Figure 2D**). The median length of distal 3’UTRs is approximately 300nt, nearly double that of genes not employing APA in their 3’UTRs. This is mostly because of the additional regulatory elements present in their sequences.

Interestingly, half of the genes that use APA (53.5%) do not possess the canonical PAS element AAUAAA in their 3’UTRs (**Main Figure 2D**). We and others have shown that APA occurs in a tissue-specific manner and, in *C. elegans,* is used in specific cellular contexts to evade miRNA-based regulatory networks in a tissue-specific manner ^13,24^. This has also been mirrored in higher eukaryotes ^22^. In this context, these AAUAAA-lacking 3’UTRs may use alternative processing mechanisms independently from the canonical CPC, which instead is ubiquitously expressed and will interfere with alternative PAS selection. However, more experiments need to be performed to identify this mechanism in more detail.

The role of the terminal adenosine and the adjacent nucleotide in the −1 position in PAS recognition and the overall cleavage reaction was unclear in our previous studies ^14^. We have now updated our *in vivo* assay and found that while the terminal adenosine plays an important role in determining the precise location of the cleavage site, the nucleotide at −1 does not (**Main Figures 4-5**). The terminal adenosine seems to modulate the location of the cleavage reaction, perhaps regulating FIPP-1 binding and/or the recruitment of the endonuclease CPSF-3. It is important to note that CPSF-3 is also implicated in the cleavage reaction of histone genes, which are not polyadenylated, but contain a terminal adenosine at their 3’ end ^38^. Perhaps CPSF-3’s predilection for cleaving at adenosine nucleotides forces this protein to be malleable and trail an adenosine nucleotide, even if it has been moved downstream of its original position in the 3’UTR. Interestingly, if this terminal adenosine is moved, the cleavage site becomes less precise ^14^. Most cleavage only occurs at nucleotides within 18 nucleotides of the cleavage site, likely because the CPC might only reach adenosine nucleotides within a certain distance from the PAS element. Interestingly, previous work on human 3’UTRs has shown that the CPC can reach adenosines further away from the PAS element, as the buffer region folds into a hairpin structure ^39^. This folding effectively shortens the length of these buffer regions, which makes it possible for the CPC to reach the cleavage site. We have identified 951 cases of 3’UTRs with buffer regions of 20 nt or more nucleotides. 125 contain canonical PAS elements (**Supplemental Table S6**). These 3’UTRs are, on average, depleted of adenosines toward the cleavage site (*data not shown*), suggesting that perhaps these longer buffer regions fold to provide a viable adenosine for CPSF-3. However, RNA folding prediction software did not reveal buffer region folding in many of these 3’UTRs (*data not shown*).

Additionally, previous *in vitro* experiments demonstrated that the cytosine preceding the terminal adenosine in human PS elements has the same effect ^40^. CPSF-3 does not interact with the nucleotide preceding the terminal adenosine during histone 3’ end processing ^41^, suggesting that this nucleotide may not have a role in the cleavage reaction during canonical pre-mRNA 3’ end processing.

Interestingly, the chemical nature of this nucleotide appears to be conserved across many eukaryotes despite its apparent lack of function. In yeast, there is an enrichment of cytosine and uracil nucleotides at −1 from the cleavage site, which is, in this organism, also an adenosine nucleotide ^42^. Several species of plants and green algae also have an enrichment of cytosine and/or uracil nucleotides at the same position ^43,44^, suggesting that eukaryotes use a pyrimidine nucleotide followed by an adenosine nucleotide at the PS site. However, the chemical nature of the nucleotide at the −1 position from the cleavage site appears to be less conserved in animals ^44^. This might indicate that this part of the PS element has been lost in animals yet remains in most other eukaryotes. Alternatively, this nucleotide might have an unknown function, such as enhancing cleavage and polyadenylation efficiency.

Notably, novel cryptic cleavage sites arise without viable adenosine at the PS site. Altering the nucleotide composition in the buffer region in our test 3’UTR had no effect on cleavage site location, mRNA transcript levels, and gene expression levels **(Main Figures 3 and 4C**), suggesting that structural constraints are not in place in this region and that the nucleotide composition of this region doesn’t have any impact on pre-mRNA processing.

We have identified a slight enrichment of uracil nucleotides at the fourth and fifth nucleotides from the PAS element (**Main Figure 3A**). These nucleotides may have a yet-uncharacterized role in the cleavage and polyadenylation reaction - perhaps as a binding site for part of the CPC during pre-mRNA 3’ end processing. Since recombinant human FIP1L1 binds to the buffer region of SV40 L3-1 RNA ^8^, we speculate that FIPP-1 may be the factor that binds to this element in the buffer region in *C. elegans* 3’UTRs.

Downstream of the cleavage sites, the G/U-rich and U-rich elements are essential for cleavage and polyadenylation in humans ^16,45^. Previous research has demonstrated that while CSTF64 binds to a UU dinucleotide within the G/U rich element, the rest of this element’s sequence can be quite variable ^17^. Notably, the functional domains of the CstF subcomplex members (which bind to this element as a dimer) are highly conserved in *C. elegans* and other metazoans ^14^.

Our study showed that the G/U-rich downstream sequence element is missing in *C. elegans*, which appears to have been replaced by a U-rich element highly abundant between +1 to +20 nucleotides. We additionally identified a bipartite U-rich element within the first 100 nucleotides downstream of the cleavage site (**Supplemental Figure S6**). Perhaps each U-rich sequence acts as a recognition motif for one of the two CSTF-2 monomers, though more work is needed to validate this hypothesis. Importantly, this element resembles those found by another group, which reanalyzed *C. elegans* EST data and identified small U-rich DSEs within the first 20 nucleotides downstream of the cleavage site ^46^.

In humans, the UGUA element is not required for the cleavage reaction to occur but, if present, acts as an enhancer for the reaction activating distal polyadenylation sites in 3’UTRs with multiple isoforms ^47^. Although the functional domains of the members of the CFIm subcomplex (which binds to the UGUA element in humans) are highly conserved between humans and worms ^14^, we were not able to either detect enrichment of the UGUA element presence or its localized distribution in 3’UTRs that use the proximal PAS site (**Supplemental Figure S7**). This was surprising since ∼25% of *C. elegans* genes use APA. Perhaps, similarly to the loss of the G/U-rich element, uracil nucleotides alone can compensate for the absence of guanosines and directly recruit the CFIm complex. Alternatively, CFIm may bind to an unknown sequence element or have a novel uncharacterized function in *C. elegans.* More research is needed to identify and characterize these 3’UTR elements in this model system.

We used miRanda ^28^ to generate updated miRNA targeting predictions for our updated 3’UTRome v3. We have prepared three miRNA targeting prediction datasets with varying levels of strictness that contain many known interactions and many yet-uncharacterized interactions that can form the basis of future research.

Our 3’UTRome v3 resource is accessible in WormBase (www.wormbase.org), a *C. elegans* genomics repository, and through our 3’UTR-centric database UTRome (www.UTRome.org) ^26^. The UTRome database is an easily accessible resource where users can visualize 3’UTR information data, download 3’UTR datasets for specific genes or in bulk, and use the genomic browser gBrowse ^48^ to display 3’UTRs and our new miRNA target predictions.

In conclusion, we here present the *C. elegans* 3’UTRome v3, which represents the most complete 3’UTR resource available in any metazoan to date and is a powerful resource for further investigations into the function of 3’UTRs in metazoans.

### Limitations of this study

We used very stringent filters when generating the *C. elegans* 3’UTRome and when doing our analysis of this data. Although these filters ensured our results were high-quality, they may have excluded some accurate 3’UTR information from this analysis. The experiments described in **Main Figures 3, 4, and 5** only used the *M03A1.3* 3’UTR. While this 3’UTR is a good representation of a typical *C. elegans* 3’UTR, it is possible that these results may not apply well to other *C. elegans* 3’UTRs.

Additionally, most of our miRNA targeting predictions have not been experimentally verified. To make our miRNA predictions dataset more specific, we narrowed the list of all the predicted miRNA targets down to those with predicted binding energies higher than 300. However, since miRanda calculates free energy based on the complementarity of the entire miRNA to the mRNA target, the strictest datasets do not contain some known interactions that involve only the seed region of the miRNA. Because of this, we have included two less strict datasets that do not have binding energy requirements.

## Acknowledgments

We thank Oliver Kask for his assistance with the Nanopore sequencing of the cloned 3’UTR isoforms. This work was funded by the National Institutes of Health 5R01GM118796 awarded to M.M.

## Author Contributions

E.M and M.M. designed the experiments. M.M. developed and executed the bioinformatic pipeline for generating the *C. elegans* 3’UTRome v3 and uploaded the 3’UTRome data to WormBase and the utrome.org website. M.M also performed the bioinformatic analyses. E.M performed the experiments shown in **Main Figures 3-5** and in **Supplemental Figure S4**. D.M. cloned the 3’UTRs shown in **Supplemental Figure S2**. N.C. manually curated PAS elements shown in this study. E.M. and M.M. prepared the main and supplemental figures and wrote the manuscript. All authors have read and approved this manuscript.

## Declaration of Interests

The authors have no conflicts of interest.

## STAR Methods

### Key Resources table

On an additional submitted file.

### Resource Availability

#### Lead contact

Requests for additional information and/or access to the materials used in this paper should be directed to the lead contact, Marco Mangone

#### Material Availability

The 3’UTR information in the *C. elegans* 3’UTRome described in this study has been uploaded to the WormBase (www.WormBase.org) and the 3’UTRome (www.UTRome.org) websites. The plasmids, primers, and transgenic *C. elegans* lines used in this study are available upon request.

## Method details

### 3’UTR mapping pipeline

Data Download: We used the SRA (Sequence Read Archive) toolkit from the National Center for Biotechnology Information (NCBI) to download raw reads from 11,533 transcriptome experiments. The data sets are listed in **Supplemental Table S1** and **Supplemental Figure S1**.

We used only datasets produced using the Illumina platform, with reads at least 100 nucleotides (nt) in length. After downloading the data, the files were unzipped and stored on our servers. We developed custom-made Perl scripts to filter the reads, extracting reads that contained either 23 consecutive adenosine nucleotides at their 3’ end or 23 consecutive thymidine nucleotides at their 5’ end. This filter resulted in 24,973,286 mappable 3’ end reads. Next, the reads were converted to FASTQ files using the FASTX-Toolkit, a tool available at http://hannonlab.cshl.edu/fastx_toolkit/. The processed reads were mapped to the WS250 release of the *C. elegans* genome using the Bowtie 2 algorithm with standard parameters^49^. A total of 7,761,642 reads (31.08%) were successfully mapped. These mapped reads were then sorted and separated based on their respective strand origin (positive or negative). Two versions of the processed data were uploaded to WormBase, a biological database for *C. elegans* research. The complete set of 3’ UTRs making up the *C. elegans* 3’UTRome v3 is provided in **Supplemental Table S2**.

### Cluster Preparation

To generate Poly(A) clusters, we followed these procedures: We retained the ID, genomic coordinates, and strand orientation of each mapped read, preserving this information for subsequent stages of our analysis. The BAM file produced by the alignment tools was first sorted and then converted into BED format using the SAMtools software^50^. To identify continuous genomic regions, we merged overlapping genomic coordinates using BEDTools software^51^. We then executed the following command: “Bedtools merge -c 1 -o count -I > tmp.cluster” to generate clusters of reads. We then separated these clusters into tiers of different stringency. We implemented several stringent filters to minimize noise in our most stringent tier, Tier 1. Clusters in this tier contained at least 50 reads. We also checked the adenosine content of the genomic DNA sequence between the end of the cluster and a site 20 nucleotides downstream of the end of the cluster. If this sequence contained more than 35% adenosine nucleotides, we excluded the cluster to exclude clusters formed by potential mispriming during the second strand synthesis in the RT reaction. Clusters were associated with the nearest gene in the same orientation. If no gene could be identified within a 1,600-nucleotide range, the cluster was disregarded. In cases where multiple 3’UTR isoforms were identified, we calculated the frequency of each isoform’s occurrence. Isoforms with a frequency of less than 30%, regardless of the number of reads forming the cluster, were omitted. We identified 22,022 3’UTR isoforms from 18,400 genes (92.8% of the genes in the WS250 release) in this tier. To detect as many 3’UTRs as possible, we loosened our filters and generated a second tier - Tier 2. To make this tier, we prepared a Bedgraph file from our mapped reads and used it to map blocks outside of known open reading frames in the genome. Then, we identified clusters in these blocks that contained at least one read and associated it with the nearest gene within 1,600 nucleotides in the same orientation. We identified an additional 1,316 3’UTR isoforms from 1,316 genes (6.7% of the genes in the WS250 release) in this tier. Lastly, we generated a third tier containing 3’UTRs that were present in the 3’UTRome v1 ^26^ but were not identified in Tiers 1 or 2. Tier 3 contained an additional 149 3’UTR isoforms from 133 genes (0.5% of the genes in the WS250 release). To visually represent our findings in **Main Figure 1E** and **Main Figure 3A**, we generated logo plots using the WebLogo 3 suite^52^.

### Identification of additional Tier 3 3’UTRs

The cloning of the 3’UTRs described in **Supplemental Figure S2** and **Supplemental Table S7** were performed as follow: total RNA from N2 wild-type *C. elegans* strain was extracted using the Zymo RNA Miniprep kit (Zymo Research). We cloned these 3’UTRs using a anchored poly-dT primer and gene specific forward DNA primers designed to encompass approximately 30 nucleotides upstream of the translation STOP codon and include the native translation STOP codon. These primers were adapted to include the necessary attB Gateway recombination elements for integration into pDONR P2r-P3 (Invitrogen). The N2 RNA was reverse transcribed using Superscript III Reverse Transcriptase and poly-dT primers under the recommended conditions (Invitrogen). The 3’UTRs of interest were amplified in a second strand reaction using Taq Polymerase (Invitrogen) with a universal anchored poly-dT primer and forward gene specific DNA primers as described above. Subsequently, we utilized the Gateway BP Clonase II Enzyme Mix from Invitrogen to insert the 3’ UTR region into pDONR P2r-P3 Gateway entry vectors. Following the recombination process, the entry vectors containing the cloned 3’ UTR regions were transformed into TOP10 competent cells (Thermo Fisher Scientific) and plated on LB agar plates containing 20 mg/µL of kanamycin. The plasmids were subsequently retrieved, and the clones were amplified using a PCR reaction using M13 forward and reverse primers and Taq Polymerase. The amplicons were then validated through Nanopore sequencing using the Ligation Sequencing Kit v14 (Oxford Nanopore Technologies). A comprehensive list of the primers used in this study can be found in **Supplemental Table S9**.

### PAS element annotation

We used Perl scripts to identify and annotate canonical and common variant PAS elements within the last 30 nucleotides of the 3’UTR isoforms identified in this study. This script could annotate PAS elements in most of the identified 3’UTRs. The remaining 266 3’UTRs were manually annotated. To identify these 3’UTRs, we searched for sequences up to 35 nt upstream of the cleavage site that closely recapitulated the chemical nature of canonical PAS elements (possessing an RRYRRR motif as well as having a third and sixth nucleotide capable of base pairing with each other).

### Maintenance of C. elegans strains

All *C. elegans* strains were cultured on NGM plates seeded with OP50-1 bacteria. All the transgenic lines produced for the experiments described in this study were maintained at 24°C and were transferred to new plates after all the bacteria on their NGM plates had been cleared.

The EG6699 strain ^53^ was maintained at 16°C. To synchronize the animals on each plate before injection, we bleached confluent plates to isolate the embryos. We washed the animals on each plate into a 15 mL Falcon tube with 15 mL DI water and pelleted for 3 minutes at 400*g*. Then, the water was aspirated, and the animals were incubated in 5 mL of a bleaching solution (0.5M NaOH, 10% bleach) for 20 minutes with gentle mixing to isolate the embryos. After bleaching, the embryos were washed twice with DI water and placed on fresh, seeded NGM plates.

### Mutagenesis of the M03A1.3 3’UTR

We used the QuikChange Site-Directed Mutagenesis Kit (Agilent) as directed to generate the *M03A1.3* 3’UTR mutants used in this study. The template constructs used were prepared in the past ^14^. The mutant M03A1.3 3’UTR shown in **Main Figure 3C** contains an additional wild-type polyadenylation site, buffer region, and PS site inserted 50 nt downstream of the original polyadenylation site in the *M03A1.3* +0A mutant 3’UTR. The mutations in the +0A, +2A, +4A, +6A, and +8A mutants are used in **Main Figure 4B**. and the mutations in the +0GA and +2TA mutants are used in **Main Figure 5B**. The products of each mutagenesis reaction were transformed into TOP10 competent cells (Invitrogen) and validated using Sanger sequencing.

### Transgenic C. elegans line preparations

We generated each transgenic *C. elegans* line from the EG6699 strain using the MosSCI approach ^53^. The injection constructs for the dual reporter lines shown in **Main Figure 3B** were generated via a Multisite LR reaction (Invitogen) using the following vectors: pCFJ150^53^, pDONR P4-P1r *myo-2p*, pDONR 221 ROG (which contains mCherry and GFP separated by a spliceable element), and either pDONR P2r-P3 *M03A1.3* 3’UTR *wt* or the +0A mutant. To generate injection constructs for the other transgenic lines described in this study, we cloned the *M03A1.3* mutants (in pDONR P2r-P3), pDONR 221 GFP, and pDONR P4-P1r *myo-2p* into CFJ150 using a Multisite LR reaction (Invitrogen). These constructs were validated using Sanger sequencing with a pCFJ150-specific M13 forward primer.

To generate the dual reporter lines shown in **Main Figure 3B**, we used injection mixes containing 25ng/µl of one of the injection constructs, 50 ng/µl pCFJ601, and 50 ng/µl pRF4^54^. To generate the rest of the transgenic lines used in this study, we used mixes containing 25ng/µl of one of the injection constructs, 50 ng/µl pCFJ601, and of 2.5 ng/µl pCFJ90^53^. These mixes were loaded onto microinjection needles, which were then mounted on to a Leica DMI300B microscope and pressurized to 28.9 psi using a Femtojet 4x (Eppendorf). Young adult EG6699 *C. elegans* were injected on 2% agarose pads covered with mineral oil on a glass coverslip. The injected animals were then placed on a fresh NGM plate seeded with OP50-1 bacteria and washed with 10 µl M9 buffer. The next day, the injected *C. elegans* were moved to individual NGM plates. The progenies of the injected animals were screened 4-7 days after injection. We screened the F1 and F2 offspring of the animals injected with the dual reporter mix for the roller phenotype. We generated and tested 5 lines of each strain, which were tested once more than 90% of the animals on each plate were transgenic. The other transgenic lines shown in the paper were screened for *unc-119* rescue and red fluorescence produced by CFJ90 expression. The transgenic F1 animals were then separated onto individual plates and allowed to grow for another 3 days. The F2 animals were then screened again. The buffer region mutant strain shown in **Main Figure 3C** was only used once a completely transgenic line had been isolated. However, the scanning insertion strains shown in **Main Figures 4** and **5** were used once more than 75% of the animals on a plate were transgenic.

### Quantification of GFP and mCherry fluorescence

Each of the transgenic *C. elegans* lines shown in this paper were imaged using a Leica DMi8 microscope. We generated 5 lines each of both dual reporter strains described in **Main Figure 3B**. To accurately quantify fluorescence levels in these lines, we imaged GFP fluorescence with 8 seconds of exposure and mCherry fluorescence with 4 seconds of exposure for each animal. We then used ImageJ to quantify the amount of fluorescence in raw images of each animal. We quantified mCherry and GFP fluorescence in the distal bulb of the pharynx of 3-4 animals from each line (totaling 21 animals from each line). We then calculated the GFP/mCherry fluorescence ratio for each animal. The fluorescence ratios of the *wt* and +0A animals were compared using a two-tailed Student’s t-test. P-values below 0.05 were considered significant.

### qPCR experiments

To quantify GFP and mCherry RNA levels in the dual reporter lines in **Main Figure 3B**, we extracted total RNA from each line (5 *wt* lines and 5 +0A lines) using the Zymo Direct-Zol RNA extraction kit (Zymo Research). This RNA was immediately reverse transcribed using Superscript III reverse transcriptase (Invitrogen). We then performed qPCR with cDNA from this reaction, SYBR Green (Invitrogen), and GFP and mCherry-specific primers under the recommended conditions. Each qPCR reaction was performed alongside a positive control reaction containing 10 pg pDONR 221 ROG instead of cDNA and a negative control reaction containing no template. GFP and mCherry RNA levels were calculated using the positive control Ct, the negative control Ct, and the sample Ct. Then, we calculated the ratio of GFP/mCherry RNA levels for each line and analyzed these values using a two-tailed Student’s t-test. P-values below 0.05 were considered significant.

### Buffer region assay

For this assay shown in **Main Figure 3C**, total RNA was extracted from different transgenic *C. elegans* line and converted to cDNA via a reverse transcription reaction using Superscript III, GFP forward primers, and anchored poly-dT primers. This cDNA was then run on a 2% agarose gel in TAE buffer alongside a 100bp-1kb DNA ladder (Bio-Rad). After electrophoresis, the gel was stained in a solution of 0.5 µg/mL ethidium bromide in TAE buffer and imaged.

### In vivo cleavage assays

For the assays shown in **Main Figure 4C** and **Main Figure 5C**, we extracted total RNA from each tested transgenic line. The RNA was converted to cDNA via reverse transcription with Superscript III Reverse transcriptase (Invitrogen) and GFP forward and poly-dT primers with attB sites at the ends. This cDNA was then cloned into pDONR P2r-P3 using a Gateway BP reaction (Invitrogen), and transformed into DH5α competent cells, and plated to isolate individual clones. Randomly selected clones were purified and sequenced using Sanger Sequencing to determine the location of cleavage in each mutant.

### miRNA targeting predictions

We used the miRanda algorithm ^28^ to generate three datasets of updated miRNA targeting predictions for the worm 3’UTRome. The strictest dataset, strict+, only contains predictions with perfect miRNA seed complementarity to its target and a binding energy score > 300. The ‘strict’ dataset contains predictions with perfect miRNA seed complementarity. Our last dataset (all) contains all the predictions produced by miRanda. The network shown in **Main Figure 6** was generated using Cytoscape^55^ and visualized using NetworkAnalyst^56^.

## Supplemental Information titles and legends

***Supplemental Figure S1*: Bioinformatic Pipeline:** We downloaded 11,533 transcriptome datasets from the SRA trace archive to identify and map 3’UTR end clusters to the nearest protein-coding genes in the correct orientation. The pipeline used to process this data was adapted from Steber *et al*. (2019). We prepared custom Perl scripts to extract reads with 23 consecutive As at the 3’ end or 23 consecutive Ts at the 5’ end. We then adapted them to fasta format and used FASTX-Toolkit (http://hannonlab.cshl.edu/fastx_toolkit/index.html) to convert them back to fastq format. The newly prepared fastq files were then mapped to the WS250 version of the *C. elegans* genome using Bowtie2 ^49^. The resulting reads were sorted and indexed. The SAM reads produced by Bowtie2 matching 100% to WS250 were extracted and utilized to prepare a new bedGraph file using BEDTools ^51^. The reads were merged, and clusters with less than 49 reads were discarded. Strict parameters for cluster identification and 3’UTR end mapping included discarding clusters with an adenosine content of less than 35% downstream of their end (to remove genomic DNA contamination errors during the cDNA preparation). The clusters were assigned to the nearest gene less than 1,600nt in the same orientation. To map 3’UTR isoforms, clusters assigned to the same gene with a density of less than 30% of the total reads were discarded. These 3’UTRs are labelled Tier 1. This tier contains the majority of the mapped 3’UTRs in this release (93% - 18,400 genes; 22,022 3’UTR isoforms). In Tier 2, the filters used to map 3’UTRs were more relaxed. We lowered the minimum number of reads in each cluster to 1, allowed a maximum of one mismatch in our mapped reads, and did not filter out clusters with more than 35% downstream adenosines. Because of all these relaxed filters, we assigned only one 3’UTR isoform to each gene (i.e., we ignored 3’UTR isoforms within this Tier). Tier 3 contains 3’UTR data extracted from Mangone *et al*., 2008, Steber *et al*., 2019, and Jan *et al*., 2011 and data from the cloned 3’UTRs described in **Supplemental Figure S2** not present in the other two tiers.

***Supplemental Figure S2*: Identification of additional Tier 3 3’UTRs** A) We attempted to clone 3’UTRs for 83 genes that were missing in our 3’UTRome dataset by reverse transcribing N2 RNA using gene-specific forward and poly-dT reverse primers. We then integrated these cDNAs into pDONR P2r-P3 using a Gateway BP reaction (Invitrogen). To validate the cloning of these 3’UTRs, we amplified each 3’UTR clone using M13 forward and reverse primers and visualized the amplicons on a 96-well agarose gel. B) M13 forward and reverse amplicons of 5 clones. C) All 83 amplicons were pooled and sequenced using Nanopore sequencing (Oxford Nanopore Technologies). We identified 3’UTR information for 28 of the 83 genes tested using this method. These 3’UTR isoforms were added to Tier 3 of our dataset.

***Supplemental Figure S3*: Examples of alternative polyadenylation events in protein-coding genes.** Screenshots from the 3’UTRome showing 3’UTR isoforms for four protein-coding genes with two 3’UTR isoforms in the 3’UTRome v3. Each panel contains the %GC content for the genomic locus shown (top), the WS250 gene model for a selected genes (second), and a model of the gene’s 3’UTR isoforms identified in this study (third), and predicted miRNA targets using our strict dataset (bottom). Notably, many miRNAs target the distal 3’UTR isoform but not the proximal 3’UTR isoform of each gene.

***Supplemental Figure S4*: Cryptic PAS usage in the +8A scanning insertion mutant.** A) The full sequence of the cloned *M03A1.3* 3’UTR used as a test 3’UTR in the *in vivo* cleavage assays in **Main Figure 4C** and **Main Figure 5C**. The end of the protein-coding sequence is highlighted in blue, the stop codon is highlighted in green, the 3’UTR is highlighted in grey, the cryptic and canonical PAS elements are underlined, the cryptic terminal adenosines are highlighted in pink, and the canonical terminal adenosines are highlighted in red. B) A model of the cryptic polyadenylation site showing the end of the coding sequence (blue), the cryptic PAS site (underlined), the stop codon (green), and the beginning of the 3’UTR (grey) (top) and trace files for both transcripts from the +8A mutant using the cryptic polyadenylation site (bottom). The PAS element at this site possesses the variant sequence TATAAA and is located at −7 nucleotides from the stop codon (purple). The cleavage site for each transcript is marked by a red dotted line.

***Supplemental Figure S5*: Characteristics of ORF-PAS 3’UTRs:** We identified 107 protein-coding genes with 3’UTR isoforms containing the PAS element upstream of the STOP codon. A) Screenshots of the genome browser from our website showing examples of these ORF-PAS 3’UTRs. The PAS elements used in these 3’UTRs are highlighted with red boxes. B) These 3’UTRs are, on average, very short, and C) Mostly possess a canonical “AAUAAA” PAS element. The complete list of ORF-PAS 3’UTRs is shown in **Supplemental Table S4.**

***Supplemental Figure S6*: U-rich element composition.** To identify potential downstream sequence elements in *C. elegans* 3’UTRs, we extracted 100nt genomic regions downstream of the PS site from 1,306 protein-coding genes with only one 3’UTR isoform containing the canonical PAS element ‘AAUAAA’ located at −16nt from the PS site that is not located in an operon. i) These regions contain a strong enrichment of uracil nucleotides at +3 to +20 (yellow box and enlarged panel). ii) Parsing these 100nt-long genomic regions with the MEME Suite ^57^ revealed an enriched symmetric bi-partite motif (E-value 7.9e-043). This motif may be used as a binding site for CSTF-2 during pre-mRNA 3’ end processing.

***Supplemental Figure S7*: UGUA motif usage.** We extracted 3’UTRs from 284 protein-coding genes that use APA, have two 3’UTR isoforms, and are not located in operons. A) We extracted the last 100nt closest to the 3 ‘ends of each isoform, removed the last 20 nucleotides, and plotted the distribution of the difference in the occurrence of the ‘UGUA’ element within these sequences. The chart is symmetrical, suggesting no enrichment of this element in both distal and proximal 3’UTRs. B) Heatmaps of ‘UGUA’ motif location between the last 100 and the last 20 nucleotides of the proximal (left) or distal (right) 3’UTR isoforms of the 284 extracted protein-coding genes. Out of these 568 3’UTRs (284 proximal and 284 distal), 276 3’UTRs possess the ‘UGUA’ element within their sequences, and their location is shown in red in the heatmaps. The ‘UGUA’ element is evenly distributed in both isoforms, suggesting that this motif is not functional in *C. elegans* 3’UTRs.

**Supplemental Table S1: Datasets used in this study.**

**Supplemental Table S2: A complete copy of the *C. elegans* 3’UTRome v3**

**Supplemental Table S3: PAS element sequences and usage in the *C. elegans* 3’UTRome (v3).**

**Supplemental Table S4: Genes that use PAS elements located within an open reading frame**

**Supplemental Table S5: miRanda targeting predictions (all, strict and strict+ predictions).**

**Supplemental Table S6: 3’UTRs with buffer regions greater than 20 nucleotides in length.**

**Supplemental Table S7: Additional Tier 3 3’UTRs.**

**Supplemental Table S8: Missing 3’UTRs.**

**Supplemental Table S9: Primers used in this study.**

## Notes

### Competing Interest Statement

The authors have declared no competing interest.

## References

1. Uncategorized References

1. Macfarlane, L.A., Murphy, P.R. (2010). MicroRNA: Biogenesis, Function and Role in Cancer. Curr Genomics 11, 537–561. doi: 10.2174/138920210793175895.

2. Mayr, C. (2019). What Are 3′ UTRs Doing? Cold Spring Harb Perspect Biol. 11. doi: 10.1101/cshperspect.a034728.

3. Boreikaite, V., Elliott, T.S., Chin, J.W., Passmore, L.Al (2022). RBBP6 activates the pre-mRNA 3’ end processing machinery in humans. Genes Dev 36, 210–224. 10.1101/gad.349223.121.

4. Schmidt, M., Kluge, F., Sandmeir, F., Kühn, U., Schäfer, P., Tüting, C., Ihling, C., Conti, E., Wahle, E. (2022). Reconstitution of 3’ end processing of mammalian pre-mRNA reveals a central role of RBBP6. Genes Dev 36, 195–209. 10.1101/gad.349217.121.

5. Sun, Y., Zhang, Y., Hamilton, K., Manley, J.L., Shi, Y., Walz, T., and Tong, L. (2018). Molecular basis for the recognition of the human AAUAAA polyadenylation signal. Proc Natl Acad Sci U S A 115, E1419-E1428. 10.1073/pnas.1718723115.

6. Clerici, M., Faini, M., Muckenfuss, L.M., Aebersold, R., and Jinek, M. (2018). Structural basis of AAUAAA polyadenylation signal recognition by the human CPSF complex. Nat Struct Mol Biol 25, 135–138. 10.1038/s41594-017-0020-6.

7. Mandel, C.R., Kaneko, S., Zhang, H., Gebauer, D., Vethantham, V., Manley, J.L., and Tong, L. (2006). Polyadenylation factor CPSF-73 is the pre-mRNA 3’-end-processing endonuclease. Nature 444, 953–956. 10.1038/nature05363.

8. Kaufmann, I., Martin, G., Friedlein, A., Langen, H., and Keller, W. (2004). Human Fip1 is a subunit of CPSF that binds to U-rich RNA elements and stimulates poly(A) polymerase. EMBO J 23, 616–626. 10.1038/sj.emboj.7600070.

9. Zhang, Y., Sun, Y., Shi, Y., Walz, T., and Tong, L. (2020). Structural Insights into the Human Pre-mRNA 3’-End Processing Machinery. Mol Cell 77, 800–809 e806. 10.1016/j.molcel.2019.11.005.

10. Mangone, M., Manoharan, A.P., Thierry-Mieg, D., Thierry-Mieg, J., Han, T., Mackowiak, S.D., Mis, E., Zegar, C., Gutwein, M.R., Khivansara, V., et al. (2010). The landscape of C. elegans 3’UTRs. Science 329, 432–435. 10.1126/science.1191244.

11. Tian, B., and Graber, J.H. (2012). Signals for pre-mRNA cleavage and polyadenylation. Wiley Interdiscip Rev RNA 3, 385–396. 10.1002/wrna.116.

12. Jan, C.H., Friedman, R.C., Ruby, J.G., and Bartel, D.P. (2011). Formation, regulation and evolution of Caenorhabditis elegans 3’UTRs. Nature 469, 97–101. 10.1038/nature09616.

13. Blazie, S.M., Babb, C., Wilky, H., Rawls, A., Park, J.G., and Mangone, M. (2015). Comparative RNA-Seq analysis reveals pervasive tissue-specific alternative polyadenylation in Caenorhabditis elegans intestine and muscles. BMC Biol 13, 4. 10.1186/s12915-015-0116-6.

14. Steber, H.S., Gallante, C., O’Brien, S., Chiu, P.L., and Mangone, M. (2019). The C. elegans 3’-UTRome v2 resource for studying mRNA cleavage and polyadenylation, 3’-UTR biology, and miRNA targeting. Genome Research.

15. Kuhn, U., and Wahle, E. (2004). Structure and function of poly(A) binding proteins. Biochim Biophys Acta 1678, 67–84. 10.1016/j.bbaexp.2004.03.008.

16. Yang, W., Hsu, P.L., Yang, F., Song, J.E., and Varani, G. (2018). Reconstitution of the CstF complex unveils a regulatory role for CstF-50 in recognition of 3’-end processing signals. Nucleic Acids Res 46, 493–503. 10.1093/nar/gkx1177.

17. Perez Canadillas, J.M., and Varani, G. (2003). Recognition of GU-rich polyadenylation regulatory elements by human CstF-64 protein. EMBO J 22, 2821–2830. 10.1093/emboj/cdg259.

18. de Vries, H., Ruegsegger, U., Hubner, W., Friedlein, A., Langen, H., and Keller, W. (2000). Human pre-mRNA cleavage factor II(m) contains homologs of yeast proteins and bridges two other cleavage factors. EMBO J 19, 5895–5904. 10.1093/emboj/19.21.5895.

19. Schafer, P., Tuting, C., Schonemann, L., Kuhn, U., Treiber, T., Treiber, N., Ihling, C., Graber, A., Keller, W., Meister, G., et al. (2018). Reconstitution of mammalian cleavage factor II involved in 3’ processing of mRNA precursors. RNA 24, 1721–1737. 10.1261/rna.068056.118.

20. Zhu, Y., Wang, X., Forouzmand, E., Jeong, J., Qiao, F., Sowd, G.A., Engelman, A.N., Xie, X., Hertel, K.J., and Shi, Y. (2018). Molecular Mechanisms for CFIm-Mediated Regulation of mRNA Alternative Polyadenylation. Mol Cell 69, 62–74 e64. 10.1016/j.molcel.2017.11.031.

21. Gruber, A.J., Zavolan, M. (2019). Alternative cleavage and polyadenylation in health and disease. Nature Reviews Genetics 20, 599–644.

22. Mayr, C., and Bartel, D.P. (2009). Widespread shortening of 3’UTRs by alternative cleavage and polyadenylation activates oncogenes in cancer cells. Cell 138, 673–684. 10.1016/j.cell.2009.06.016.

23. Curinha, A., Brax, S.O., Pereira-Castro, I., Cruz, A., Moreira, A. (2014). Implications of polyadenylation in health and disease. Nucleus 5, 508–519. 10.4161/nucl.36360.

24. Blazie, S.M., Geissel, H.C., Wilky, H., Joshi, R., Newbern, J., and Mangone, M. (2017). Alternative Polyadenylation Directs Tissue-Specific miRNA Targeting in Caenorhabditis elegans Somatic Tissues. Genetics 206, 757–774. 10.1534/genetics.116.196774.

25. Sanfilippo, P., Miura, P., and Lai, E.C. (2017). Genome-wide profiling of the 3’ ends of polyadenylated RNAs. Methods 126, 86–94. 10.1016/j.ymeth.2017.06.003.

26. Mangone, M., Macmenamin, P., Zegar, C., Piano, F., and Gunsalus, K.C. (2008). UTRome.org: a platform for 3’UTR biology in C. elegans. Nucleic Acids Res 36, D57–62. 10.1093/nar/gkm946.

27. Schorr, A.L., Mejia, A.F., Miranda, M.Y., and Mangone, M. (2023). An updated C. elegans nuclear body muscle transcriptome for studies in muscle formation and function. Skelet Muscle 13, 4. 10.1186/s13395-023-00314-2.

28. Enright, A.J., John, B., Gaul, U., Tuschl, T., Sander, C., Marks, D.S. (2003). MicroRNA targets in Drosophila. Genome Biol 5. 10.1186/gb-2003-5-1-r1.

29. Abrahante, J.E., Daul, A.L., Li, M., Volk, M.L., Tennessen, J.M., Miller, E.A., Rougvie, A.E. (2003). The Caenorhabditis elegans hunchback-like gene lin-57/hbl-1 controls developmental time and is regulated by microRNAs. Dev Cell 4, 625–637. 10.1016/s1534-5807(03)00127-8.

30. Grosshans, H., Johnson, T., Reinert, K.L., Gerstein, M., Slack, F.J. (2005). The temporal patterning microRNA let-7 regulates several transcription factors at the larval to adult transition in C. elegans. Dev Cell 8, 321–330. 10.1016/j.devcel.2004.12.019.

31. Johnston, R.J., Hobert, O. (2003). A microRNA controlling left/right neuronal asymmetry in Caenorhabditis elegans. Nature 426, 845–849. 10.1038/nature02255.

32. Ecsedi, M., Rausch, M., Großhans, H. The let-7 microRNA directs vulval development through a single target. Dev Cell 32, 335–344. 10.1016/j.devcel.2014.12.018.

33. Aeschimann, F., Neagu, A., Rausch, M., Großhans, H. (2019). let-7 coordinates the transition to adulthood through a single primary and four secondary targets. Life Sci Alliance 2. 10.26508/lsa.201900335.

34. Johnson, S.M., Grosshans, H., Shingara, J., Byrom, M., Jarvis, R., Cheng, A., Labourier, E., Reinert, K.L., Brown, D., Slack, F.J. (2005). RAS is regulated by the let-7 microRNA family. Cell 120, 635–641.

35. Moss, E.G., Lee, R.C., Ambros, V. (1997). The cold shock domain protein LIN-28 controls developmental timing in C. elegans and is regulated by the lin-4 RNA. Cell 88, 637–646. 0.1016/s0092-8674(00)81906-6.

36. Proudfoot, N.J., and Brownlee, G.G. (1976). 3’ non-coding region sequences in eukaryotic messenger RNA. Nature 263, 211–214. 10.1038/263211a0.

37. Fitzgerald, M., Shenk, T. (1981). The sequence 5’-AAUAAA-3’forms parts of the recognition site for polyadenylation of late SV40 mRNAs. Cell 24, 251–260. 10.1016/0092-8674(81)90521-3.

38. Dominski, Z., and Marzluff, W.F. (2007). Formation of the 3’ end of histone mRNA: getting closer to the end. Gene 396, 373–390. 10.1016/j.gene.2007.04.021.

39. Wu, X., Bartel, D.P. (2017). Widespread Influence of 3’-End Structures on Mammalian mRNA Processing and Stability. Cell 169, 905–917. 10.1016/j.cell.2017.04.036.

40. Chen, F., MacDonald, C.C., and Wilusz, J. (1995). Cleavage site determinants in the mammalian polyadenylation signal. Nucleic Acids Res 23, 2614–2620.

41. Sun, Y., Zhang, Y., Aik, W.S., Yang, X.C., Marzluff, W.F., Walz, T., Dominski, Z., Tong, L. (2020). Structure of an active human histone pre-mRNA 3’-end processing machinery. Science 367, 700–703. 10.1126/science.aaz7758.

42. Dichtl, B., Keller, W. (2001). Recognition of polyadenylation sites in yeast preLmRNAs by cleavage and polyadenylation factor. EMBO J 20. 10.1093/emboj/20.12.3197.

43. Wodniok, S., Simon, A., Glöckner, G., Becker, B. (2007). Gain and loss of polyadenylation signals during evolution of green algae. BMC Evol Biol. 7. 10.1186/1471-2148-7-65.

44. Li, X.Q., Du, D. (2014). Motif types, motif locations and base composition patterns around the RNA polyadenylation site in microorganisms, plants and animals. BMC Evol Biol. 14. 10.1186/s12862-014-0162-7.

45. McDevitt, M.A., Hart, R.P., Wong, W.W., Nevins, J.R. (1986). Sequences capable of restoring poly(A) site function define two distinct downstream elements. EMBO J 5, 2907–2913. 10.1002/j.1460-2075.1986.tb04586.x.

46. Salisbury, J., Hutchison, K.W., Graber, J.H. (2006). A multispecies comparison of the metazoan 3’-processing downstream elements and the CstF-64 RNA recognition motif. BMC Genomics 7. 10.1186/1471-2164-7-55.

47. Ghosh, S., Ataman, M., Bak, M., Börsch, A., Schmidt, A., Buczak, K., Martin, G., Dimitriades, B., Herrmann, C.J., Kanitz, A., Zavolan, M. (2022). CFIm-mediated alternative polyadenylation remodels cellular signaling and miRNA biogenesis. Nucleic Acids Res 50, 3096–3114. 10.1093/nar/gkac114.

48. Stein, L.D., Mungall, C., Shu, S., Caudy, M., Mangone, M., Day, A., Nickerson, E., Stajich, J.E., Harris, T.W., Arva, A., and Lewis, S. (2002). The generic genome browser: a building block for a model organism system database. Genome Res 12, 1599–1610. 10.1101/gr.403602.

49. Langmead, B., and Salzberg, S.L. (2012). Fast gapped-read alignment with Bowtie 2. Nat Methods 9, 357–359. 10.1038/nmeth.1923.

50. Li, H., Handsaker, B., Wysoker, A., Fennell, T., Ruan, J., Homer, N., Marth, G., Abecasis, G., Durbin, R., and Genome Project Data Processing, S. (2009). The Sequence Alignment/Map format and SAMtools. Bioinformatics 25, 2078–2079. 10.1093/bioinformatics/btp352.

51. Quinlan, A.R., and Hall, I.M. (2010). BEDTools: a flexible suite of utilities for comparing genomic features. Bioinformatics 26, 841–842. 10.1093/bioinformatics/btq033.

52. Crooks, G.E., Hon, G., Chandonia, J.M., and Brenner, S.E. (2004). WebLogo: a sequence logo generator. Genome Res 14, 1188–1190. 10.1101/gr.849004.

53. Frokjaer-Jensen, C., Davis, M.W., Hopkins, C.E., Newman, B.J., Thummel, J.M., Olesen, S.P., Grunnet, M., and Jorgensen, E.M. (2008). Single-copy insertion of transgenes in Caenorhabditis elegans. Nat Genet 40, 1375–1383. 10.1038/ng.248.

54. Mello, C.C., Kramer, J.M., Stinchcomb, D., and Ambros, V. (1991). Efficient gene transfer in C.elegans: extrachromosomal maintenance and integration of transforming sequences. EMBO J 10, 3959–3970.

55. Shannon, P., Markiel, A., Ozier, O., Baliga, N.S., Wang, J.T., Ramage, D., Amin, N., Schwikowski, B., Ideker, T. (2003). Cytoscape: a software environment for integrated models of biomolecular interaction networks. Genome Res 13, 2498–2504. 10.1101/gr.1239303.

56. Zhou, G., Soufan, O., Ewald, J., Hancock, R.E.W., Basu, N. and Xia, J. (2019). NetworkAnalyst 3.0: a visual analytics platform for comprehensive gene expression profiling and meta-analysis. Nucleic Acids Res 47, W234–W241. 10.1093/nar/gkz240.

57. Bailey, T.L., Boden, M., Buske, F.A., Frith, M., Grant, C.E., Clementi, L., Ren, J., Li, W.W., and Noble, W.S. (2009). MEME SUITE: tools for motif discovery and searching. Nucleic Acids Res 37, W202–208. 10.1093/nar/gkp335.

