## Supplemental Figures for "A Comprehensive Analysis of 3’UTRs in *Caenorhabditis elegans*"

Additional File 1 for:

**This PDF includes:**

**Supplemental Figs. S1-S7**

**Fig. S7: UGUA motif usage** .....................…………………….……...10

**Supplemental Fig. S1**

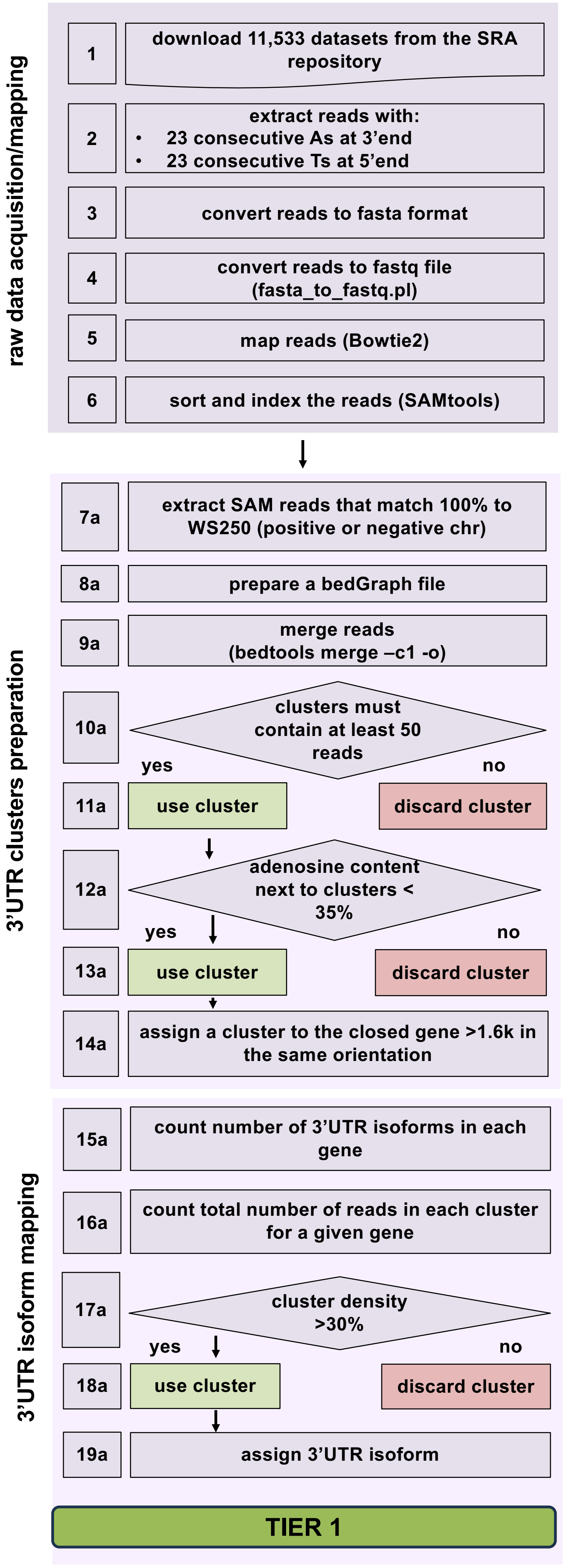

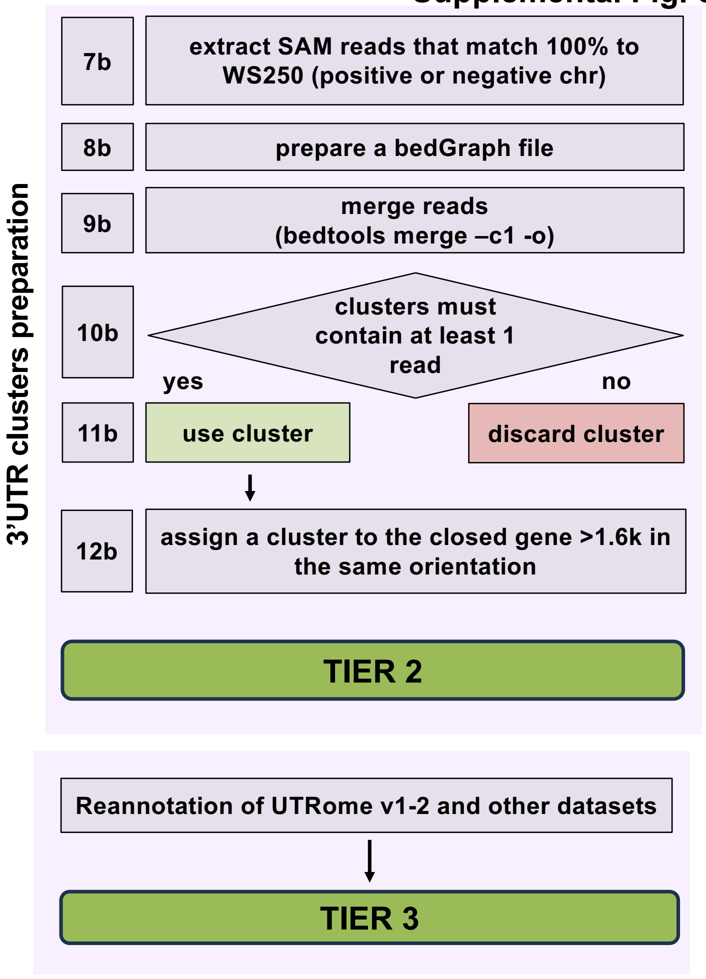

**Supplemental Fig. S1: Bioinformatic pipeline:** We downloaded 11,533 transcriptome datasets from the SRA trace archive to identify and map 3’UTR end clusters to the nearest protein-coding genes in the correct orientation. The pipeline used to process this data was adapted from Steber *et al*. (2019).
We prepared custom Perl scripts to extract reads with 23 consecutive As at the 3’ end or 23 consecutive Ts at the 5’ end. We then adapted them to fasta format and used FASTX-Toolkit (http://hannonlab.cshl.edu/fastx_toolkit/index.html) to convert them back to fastq format.
The newly prepared fastq files were then mapped to the WS250 version of the *C. elegans* genome using Bowtie2 [49]. The resulting reads were sorted and indexed. The SAM reads produced by Bowtie2 matching 100% to WS250 were extracted and utilized to prepare a new bedGraph file using BEDTools [51]. (*continued on next page*)

(*continued from previous page*) The reads were merged, and clusters with less than 49 reads were discarded. Strict parameters for cluster identification and 3'UTR end mapping included discarding clusters with an adenosine content of less than 35% downstream of their end (to remove genomic DNA contamination errors during the cDNA preparation). The clusters were assigned to the nearest gene less than 1,600nt in the same orientation. To map 3'UTR isoforms, clusters assigned to the same gene with a density of less than 30% of the total reads were discarded. These 3'UTRs are labelled Tier 1. This tier contains the majority of the mapped 3'UTRs in this release (93% - 18,400 genes; 22,022 3'UTR isoforms). In Tier 2, the filters used to map 3'UTRs were more relaxed. We lowered the minimum number of reads in each cluster to 1, allowed a maximum of one mismatch in our mapped reads, and did not filter out clusters with more than 35% downstream adenosines. Because of all these relaxed filters, we assigned only one 3'UTR isoform to each gene (i.e., we ignored 3'UTR isoforms within this Tier). Tier 3 contains 3'UTR data extracted from Mangone *et al*., 2008, Steber *et al*., 2019, and Jan *et al*., 2011 not present in the other two tiers.

**A**

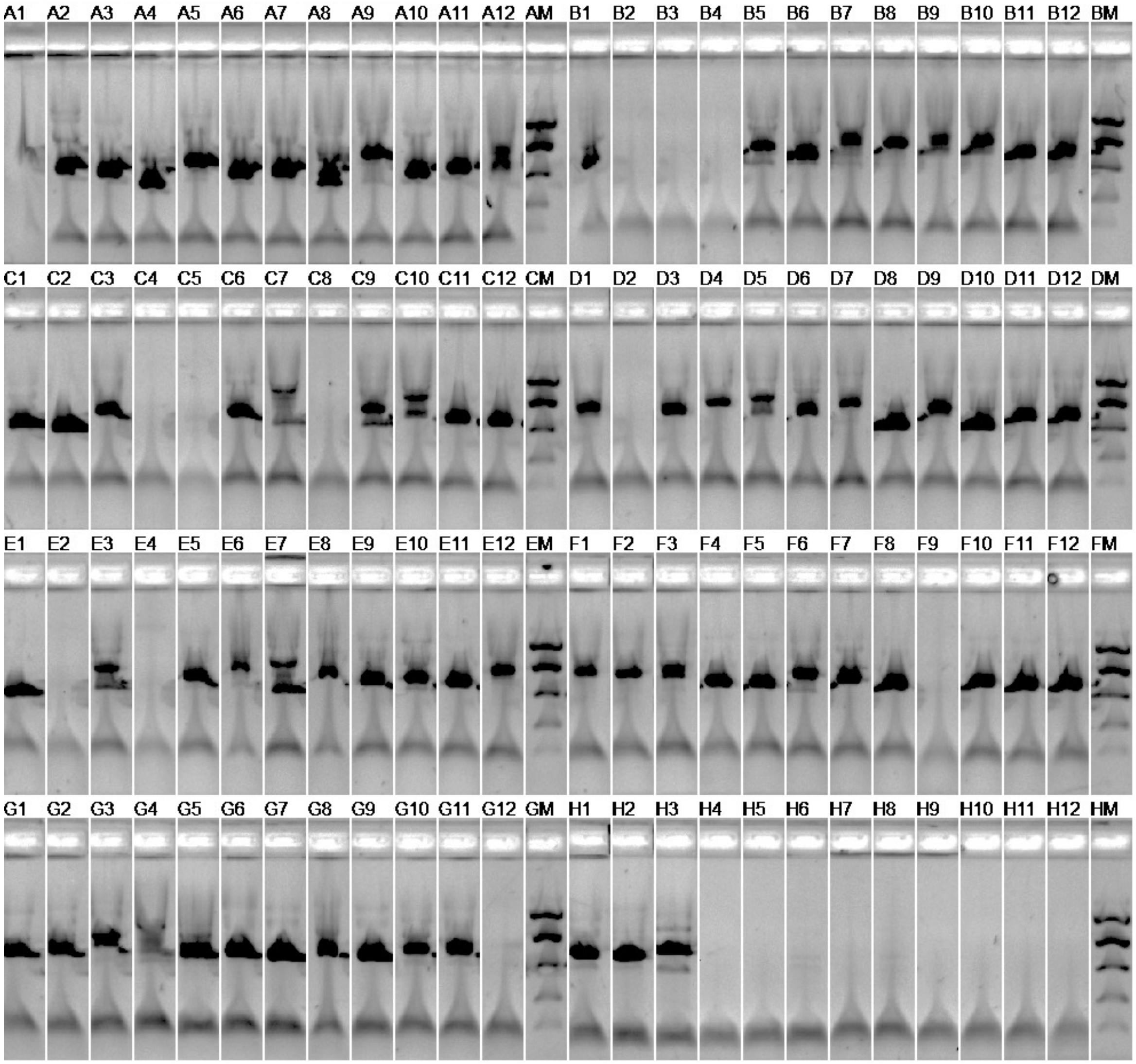

**B**

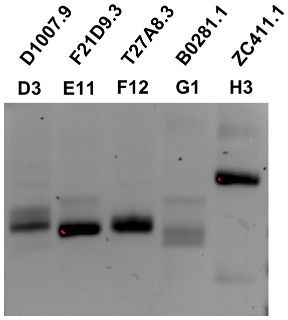

**C**

| | Cloned | Identified | Missing |
|---|---|---|---|
| genes | 83 | 25 (30%) | 55 (70%) |
| 3'UTR isoforms | - | 25 | - |

**Supplemental Fig. S2: Identification of additional Tier 3 3'UTRs:** A) We attempted to clone 3'UTRs for 83 genes that were missing in our 3'UTRome dataset by reverse transcribing N2 RNA using gene-specific forward and poly-dT reverse primers. We then integrated these cDNAs into pDONR P2r-P3 using a Gateway BP reaction (Invitrogen). To validate the cloning of these 3'UTRs, we amplified each 3'UTR clone using M13 forward and reverse primers and visualized the amplicons on a 96-well agarose gel. B) M13 forward and reverse amplicons of 5 clones. C) All 83 amplicons were pooled and sequenced using Nanopore sequencing (Oxford Nanopore Technologies). We identified 3'UTR information for 28 of the 83 genes tested using this method. These 3'UTR isoforms were added to Tier 3 of our dataset.

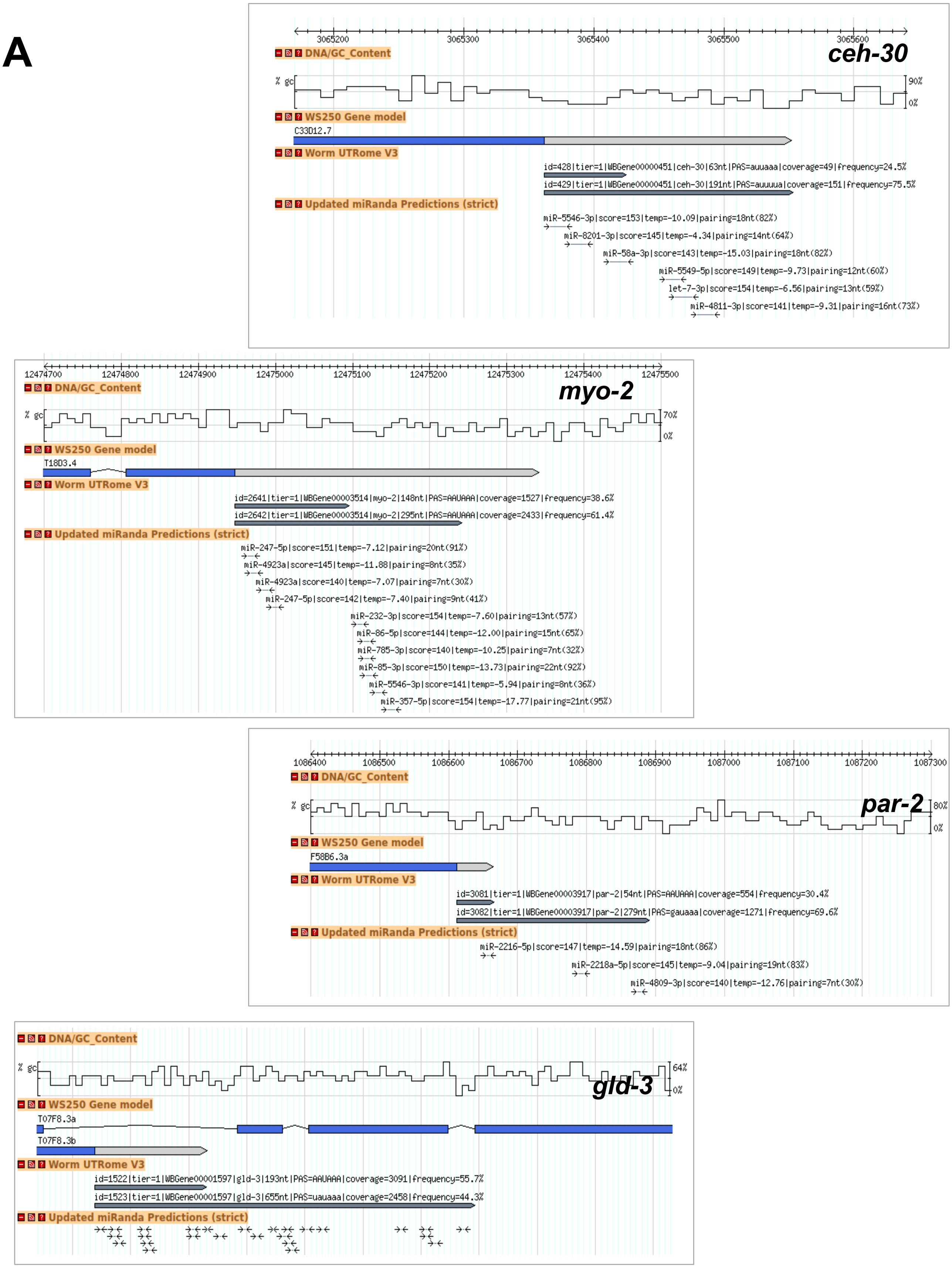

**Supplemental Fig. S3: Examples of alternative polyadenylation events in protein coding genes.** Screenshots from the 3'UTRome showing 3'UTR isoforms for four protein-coding genes with two 3'UTR isoforms in the 3'UTRome v3. Each panel contains the %GC content for the genomic locus shown (top), the WS250 gene model for a selected genes (second), and a model of the gene's 3'UTR isoforms identified in this study (third), and predicted miRNA targets using our strict dataset (bottom). Notably, many miRNAs target the distal 3'UTR isoform but not the proximal 3'UTR isoform of each gene.

## A

TGAAAGGACCTGCAGTGTTTTGGGCGATTGGAGTATTCTTCTGCATTGCTGTTGCGT
TGTCACTTCTTGTCGTCAATGGATATAAAAATGTATAATTATTAATGGAATTTTGGA
ATCTCATCTAATTTATTGATTTTATTGAATACGGGTAGTTTCTGATAATTACTTTGC
ATTGTAAAAAAACAAACTTTGTATGAATAAACATATTGAACATCTAAGTGCTTGCGT
TTTTTTAAACTCAACTTTGGTTGCGCATATCTTGGCTCTCTTTAGTTTTTATTAAAA
AATGTCAACTACAGAAATATTTGTCAATTCTTTTTCTATAGTTTTGTAGTTAATT

## B

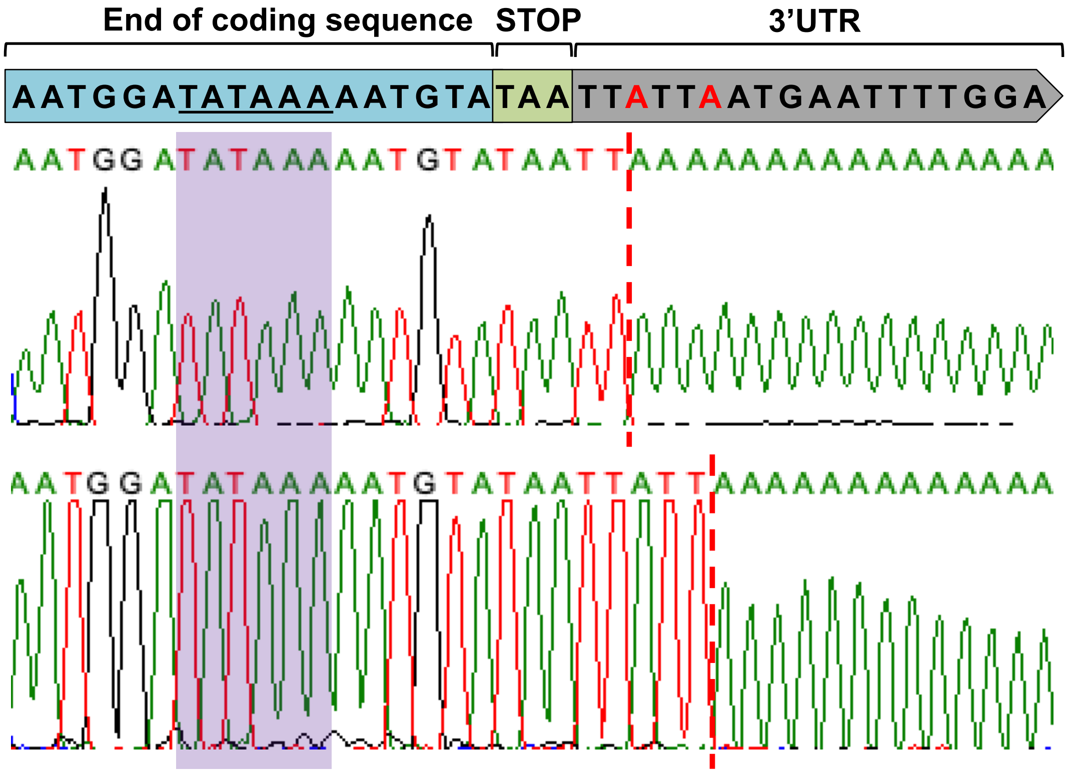

**Supplemental Fig. S4: Cryptic PAS element usage in the +8A scanning insertion mutant:** A) The full sequence of the cloned *M03A1.3* 3'UTR used as a test 3'UTR in the *in vivo* cleavage assays. The end of the protein-coding sequence is highlighted in blue, the stop codon is highlighted in green, the 3'UTR is highlighted in grey, the cryptic and canonical PAS elements are underlined, the cryptic terminal adenosines are highlighted in pink, and the canonical terminal adenosines are highlighted in red. B) A model of the cryptic polyadenylation site showing the end of the coding sequence (blue), the cryptic PAS site (underlined), the stop codon (green), and the beginning of the 3'UTR (grey) (top) and trace files for both transcripts from the +8A mutant using the cryptic polyadenylation site (bottom). The PAS element at this site possesses the variant sequence TATAAA and is located at -7 nucleotides from the stop codon (purple). The cleavage site for each transcript is marked by a red dotted line.

A

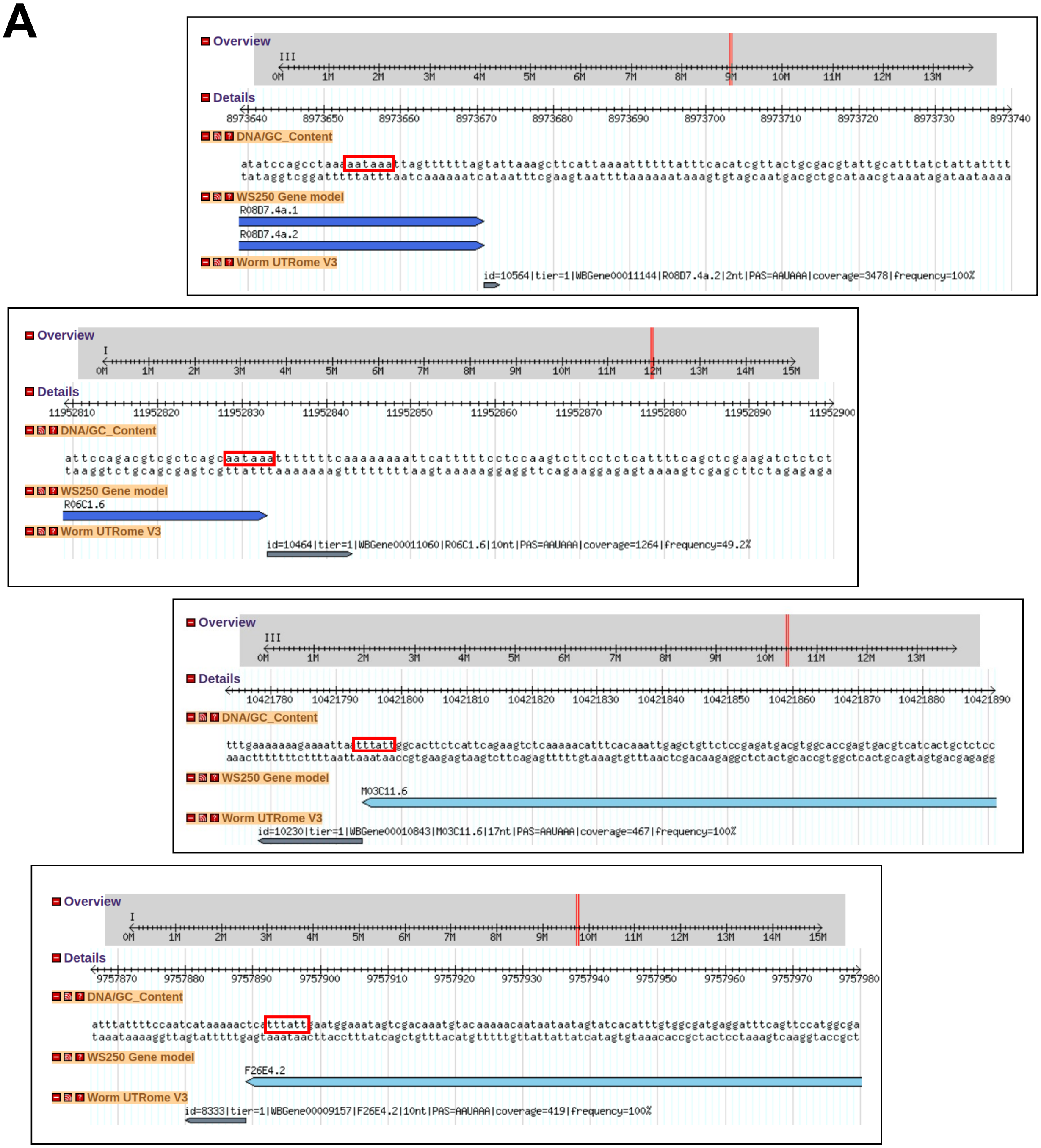

B

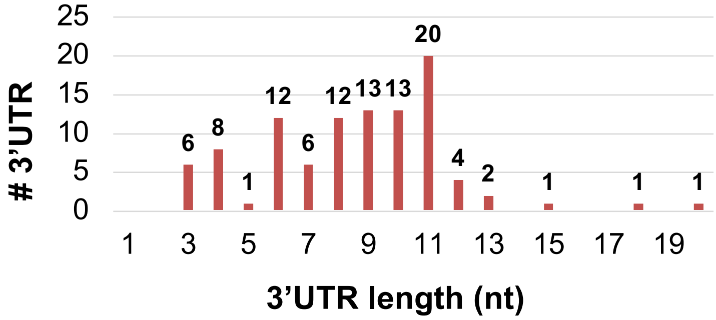

C

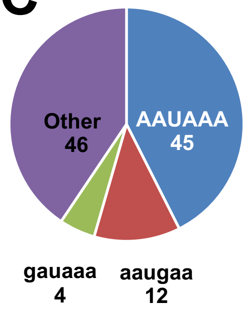

**Supplemental Fig. S5: Characteristics of ORF-PAS 3'UTRs:** We identified 107 protein-coding genes with 3'UTR isoforms containing the PAS element upstream of the STOP codon. A) Screenshots of the genome browser from our website showing examples of these ORF-PAS 3'UTRs. The PAS elements used in these 3'UTRs are highlighted with red boxes. B) These 3'UTRs are, on average, very short, and C) Mostly possess a canonical "AAUAAA" PAS element. The complete list of ORF-PAS 3'UTRs is shown in **Supplemental Table S4.**

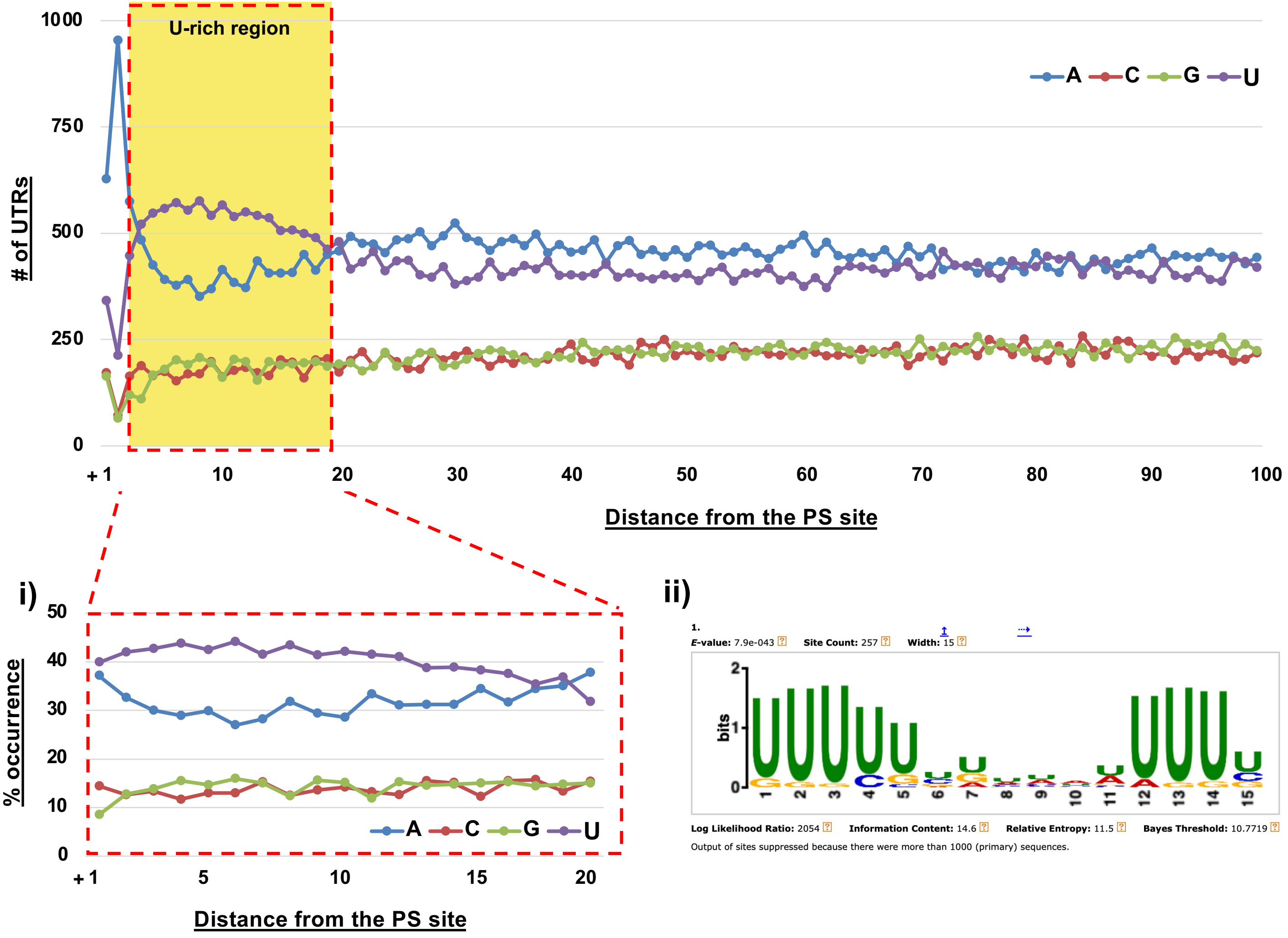

**Supplemental Fig. S6: U-rich element composition:** To identify potential downstream sequence elements in *C. elegans* 3'UTRs, we extracted 100nt genomic regions downstream of the PS site from 1,306 protein-coding genes with only one 3'UTR isoform containing the canonical PAS element 'AAUAAA' located at -16nt from the PS site that is not located in an operon. i) These regions contain a strong enrichment of uracil nucleotides at +3 to +20 (yellow box and enlarged panel). ii) Parsing these 100nt-long genomic regions with the MEME Suite [57] revealed an enriched symmetric bi-partite motif (E-value 7.9e-043). This motif may be used as a binding site for CSTF-2 during pre-mRNA 3' end processing.

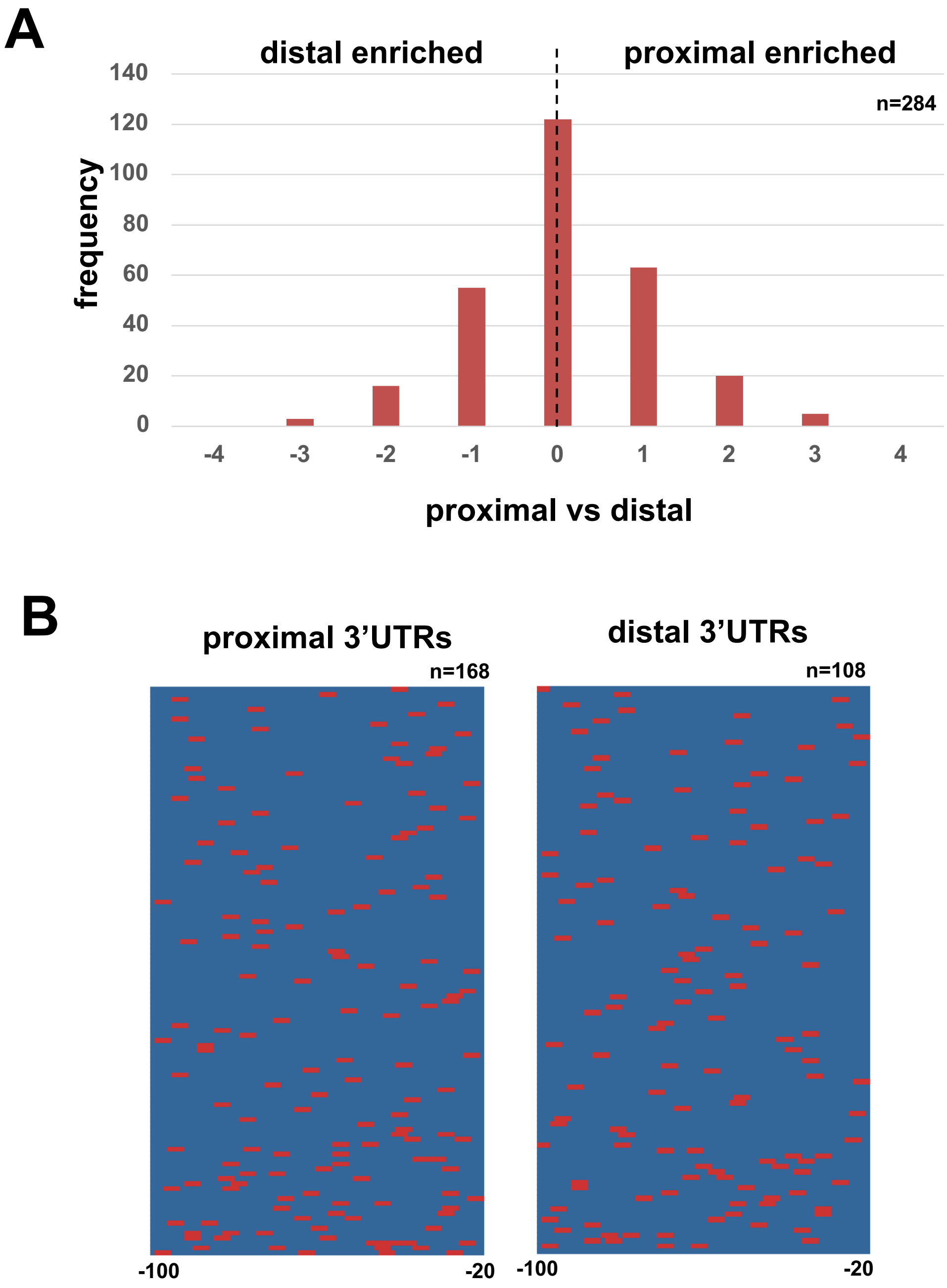

**Supplemental Fig. S7: UGUA motif usage:** We extracted 3'UTRs from 284 protein-coding genes that use APA, have two 3'UTR isoforms, and are not located in operons. A) We extracted the last 100nt closest to the 3 'ends of each isoform, removed the last 20 nucleotides, and plotted the distribution of the difference in the occurrence of the 'UGUA' element within these sequences. The chart is symmetrical, suggesting no enrichment of this element in both distal and proximal 3'UTRs. B) Heatmaps of 'UGUA' motif location between the last 100 and the last 20 nucleotides of the proximal (left) or distal (right) 3'UTR isoforms of the 284 extracted protein coding genes. Out of these 568 3'UTRs (284 proximal and 284 distal), 276 3'UTRs possess the 'UGUA' element within their sequences, and their location is shown in red in the heatmaps. The 'UGUA' element is evenly distributed in both isoforms, suggesting that this motif is not functional in *C. elegans* 3'UTRs.

### References

1. Steber, H.S., Gallante, C., O'Brien, S., Chiu, P.L., and Mangone, M. (2019). The C. elegans 3'-UTRome v2 resource for studying mRNA cleavage and polyadenylation, 3'-UTR biology, and miRNA targeting. Genome Research

2. Langmead, B., and Salzberg, S.L. (2012). Fast gapped-read alignment with Bowtie 2. Nat Methods *9*, 357-359. 10.1038/nmeth.1923.

3. Quinlan, A.R., and Hall, I.M. (2010). BEDTools: a flexible suite of utilities for comparing genomic features. Bioinformatics *26*, 841-842. 10.1093/bioinformatics/btq033.

4. Mangone, M., Manoharan, A.P., Thierry-Mieg, D., Thierry-Mieg, J., Han, T., Mackowiak, S.D., Mis, E., Zegar, C., Gutwein, M.R., Khivansara, V., et al. (2010). The landscape of C. elegans 3'UTRs. Science *329*, 432-435. 10.1126/science.1191244.

5. Jan, C.H., Friedman, R.C., Ruby, J.G., and Bartel, D.P. (2011). Formation, regulation and evolution of Caenorhabditis elegans 3'UTRs. Nature *469*, 97-101. 10.1038/nature09616.
